# Infection and antibiotic-associated changes in the fecal microbiota of *C. rodentium* ϕ*stx2_dact_*-infected C57BL/6 mice

**DOI:** 10.1101/2023.12.21.572832

**Authors:** Sabrina Mühlen, Ann Kathrin Heroven, Bettina Elxnat, Silke Kahl, Dietmar H. Pieper, Petra Dersch

## Abstract

Enterohemorrhagic *E. coli* causes watery to bloody diarrhea, which may progress to hemorrhagic colitis and hemolytic-uremic syndrome. While early studies suggested that antibiotic treatment may worsen the pathology of an EHEC infection, recent work has shown that certain non-Shiga toxin-inducing antibiotics avert disease progression. Unfortunately, both intestinal bacterial infections and antibiotic treatment are associated with dysbiosis. This can alleviate colonization resistance, facilitate secondary infections, and potentially lead to more severe illness. To address the consequences in the context of an EHEC infection, we used the established mouse infection model organism *C. rodentium* ϕ*stx2_dact_* and monitored changes in fecal microbiota composition during infection and antibiotic treatment. *C. rodentium* ϕ*stx2_dact_* infection resulted in minor changes compared to antibiotic treatment. The infection caused clear alterations in the microbial community, leading mainly to a reduction of Muribaculaceae and a transient increase in Enterobacteriaceae distinct from *Citrobacter*. Antibiotic treatments of the infection resulted in marked and distinct variations in microbiota composition, diversity, and dispersion. Enrofloxacin and trimethoprim/sulfamethoxazole, which did not prevent Stx-mediated organ damage, had the least disruptive effects on the intestinal microbiota, while kanamycin and tetracycline, which rapidly cleared the infection, caused a severe reduction in diversity. Kanamycin treatment resulted in the depletion of all but Bacteroidetes genera, whereas tetracycline effects on Clostridia were less severe. Together, these data highlight the need to address the impact of individual antibiotics in the clinical care of life-threatening infections and consider microbiota-regenerating therapies.

**IMPORTANCE:** Understanding the impact of antibiotic treatment on enterohemorrhagic *E. coli* (EHEC) infections is crucial for appropriate clinical care. While discouraged by early studies, recent findings suggest certain antibiotics can impede disease progression. Here, we investigated the impact of individual antibiotics on the fecal microbiota in the context of an established EHEC mouse model using *C. rodentium* ϕ*stx2_dact_*. The infection caused significant variations in the microbiota, leading to a transient increase in Enterobacteriaceae distinct from *Citrobacter*. However, these effects were minor compared to those observed for antibiotic treatments. Indeed, antibiotics that most efficiently cleared the infection also had the most detrimental effect on the fecal microbiota, causing a substantial reduction in microbial diversity. Conversely, antibiotics showing adverse effects or incomplete bacterial clearance had a reduced impact on microbiota composition and diversity. Taken together, our findings emphasize the delicate balance required to weigh the harmful effects of infection and antibiosis in treatment.

## INTRODUCTION

Infections with bacterial pathogens are still among the most common causes of death globally (1). With the discovery of penicillin in 1928 (2), mortality rates associated with bacterial infections decreased significantly (3). However, research has also shown that the use of antibiotics increases the development of antibiotic-resistant pathogens (1, 3, 4), can enhance susceptibility to other intestinal pathogens (5, 6), or can increase the risk of complications, as in case of infections with enterohemorrhagic *E. coli* (EHEC) (7–9).

Infections with enterohaemorrhagic *E. coli* cause bloody diarrhea and can progress to hemorrhagic colitis and hemolytic-uremic syndrome (HUS) in approximately 10 to 15% of cases (10). Children are most susceptible to infection, but adults, especially the elderly, are also affected. Shiga toxin (Stx) is considered the major virulence factor of EHEC (7, 10–12) and is responsible for its increased virulence compared to enteropathogenic *E. coli* (EPEC) (11, 12). To date, the treatment options are still limited and primarily consist of supportive therapies as it has been suggested that antibiotic treatment may cause a worsening of the disease due to increased production and release of Shiga toxin (8, 9). Several studies have investigated the ability of different antibiotics to induce Shiga toxin production *in vitro* and *in vivo* (7–9, 13), but the results obtained were inconclusive.

Due to the high host specificity of the pathogen, no proper mouse model was previously available to study EHEC infections *in vivo*. While *C. rodentium* is a suitable model for EPEC infections (14), it does not encode Shiga toxin. In 2012, a *C. rodentium* strain was generated, which encodes the Stx2_dact_ phage (*C. rodentium* ϕ*stx2_dact_*), mimicking EHEC and providing a model to study its pathogenesis *in vivo* (15). Using this mouse model, we demonstrated that the application of *stx*-inducing antibiotics resulted in weight loss and kidney damage despite the clearance of infection (16). However, several non-*stx*-inducing antibiotics cleared the bacterial infection without causing Stx-mediated pathology, suggesting that these antibiotics might be useful for treating EHEC infections (16).

Unfortunately, many antibiotics are known to alter the composition and richness of the microbiota, resulting in dysbiosis, which can be unfavorable for overall health (17, 18). The intestinal microbiota of mammals has a significant impact on metabolic and nutritional activities (19, 20), host immune responses (e.g., immune cell population, cytokine patterns (21–23), and behavioral patterns (24). An intact microbiota provides beneficial biological functions by producing metabolic products such as vitamins and short-chain fatty acids (SCFAs) (25). It is thus not surprising that antibiotic-mediated changes of the microbiota can be associated with intestinal disorders such as inflammatory bowel disease, diarrhea, and colitis, as well as extraintestinal and systemic disorders, including metabolic diseases (diabetes), autoimmune responses (rheumatoid arthritis), allergies, asthma, obesity, and even neoplastic and neurodegenerative diseases (19, 26–28).

Another consequence of antibiotic-triggered dysbiosis is the loss of colonization resistance against pathogens (26). A prominent example is treatment with antibiotics such as clindamycin and vancomycin, which leads to increased and long-lasting susceptibility to *Clostridioides difficile* and allows expansion and dense colonization of resistant *Enterococcus* and *Klebsiella pneumoniae* strains (5, 6). Recently, it was also shown that colonization of mice with *C. rodentium* and *C. rodentium* ϕ*stx2_dact_-*mediated pathology varied greatly, depending on the intestinal microbiota composition and the production of SCFAs (29).

Although several studies have investigated the changes in the murine gastrointestinal microbiota composition in response to antibiotic treatment (30–37), there is little information on the individual influence of different classes of antibiotics on the microbiota composition in a murine infection model. Additionally, there are no studies that assess the impact of infection on the fecal microbiota in the presence and absence of antibiotics with Shiga toxin-producing *C. rodentium*. For this reason, we studied the impact of *C. rodentium* ϕ*stx2_dact_* infection and antibiotic treatment on the fecal microbiota composition as a measure of intestinal microbiota disruption. We found that *C. rodentium* ϕ*stx2_dact_* infection caused significant alterations in the composition of the murine fecal microbiota. These changes were, however, minor in magnitude and direction compared to the effects of antibiotics, which resulted in tremendous and highly diverse antibiotic-specific changes in microbiota structure and diversity. Knowledge of the destructive effect on potentially beneficial commensals triggered by a specific class of antibiotics may be helpful for clinical applications, such as treating enteric infections.

## RESULTS

Antibiotics are commonly used to treat bacterial infections. However, as some antibiotics are known to induce the expression and secretion of Shiga toxin, they are generally not suggested as a treatment strategy for EHEC infections (7, 8). In an earlier study, we systematically analyzed the effect of antibiotics from different classes on Shiga toxin-mediated disease using *C. rodentium* ϕ*stx2_dact_*. In this study, mice were orally infected with *C. rodentium* ϕ*stx2_dact_*. Starting from day 4 post-infection, mice were daily treated with five antibiotics of different classes (enrofloxacin – Enf, kanamycin – Kan, rifampicin – Rif, tetracycline – Tet; and trimethoprim/sulfamethoxazole – T/S). Throughout the study, fecal samples were collected to allow the analysis of the microbiota composition on days 0 (before infection), 4, 6, and 12 post-infection (**Fig. 1**). These time points were chosen as *C. rodentium* initiates colonization of the colon on day 4, reaches a maximal uniform distribution along the entire colonic mucosa on day 6 and starts to decline on day 12 post-infection (38). In total, we analyzed 410 samples of 125 mice belonging to seven different treatment groups (uninfected (UI), infected untreated (UT), and infected, antibiotic-treated mice (Enf, Kan, Rif, Tet, T/S) at the four time-points (**Fig. 1**) by 16S rRNA gene sequencing assessing the effects of *C. rodentium* ϕ*stx2_dact_* infection as well as short-and long-term antibiotics treatment in C57BL/6Rj mice **(Table S1).** In total, 7,654,419 bacterial 16S rDNA sequence counts were obtained with a mean of 18,669 ± 8,447 counts per sample.

**Fig. 1.**
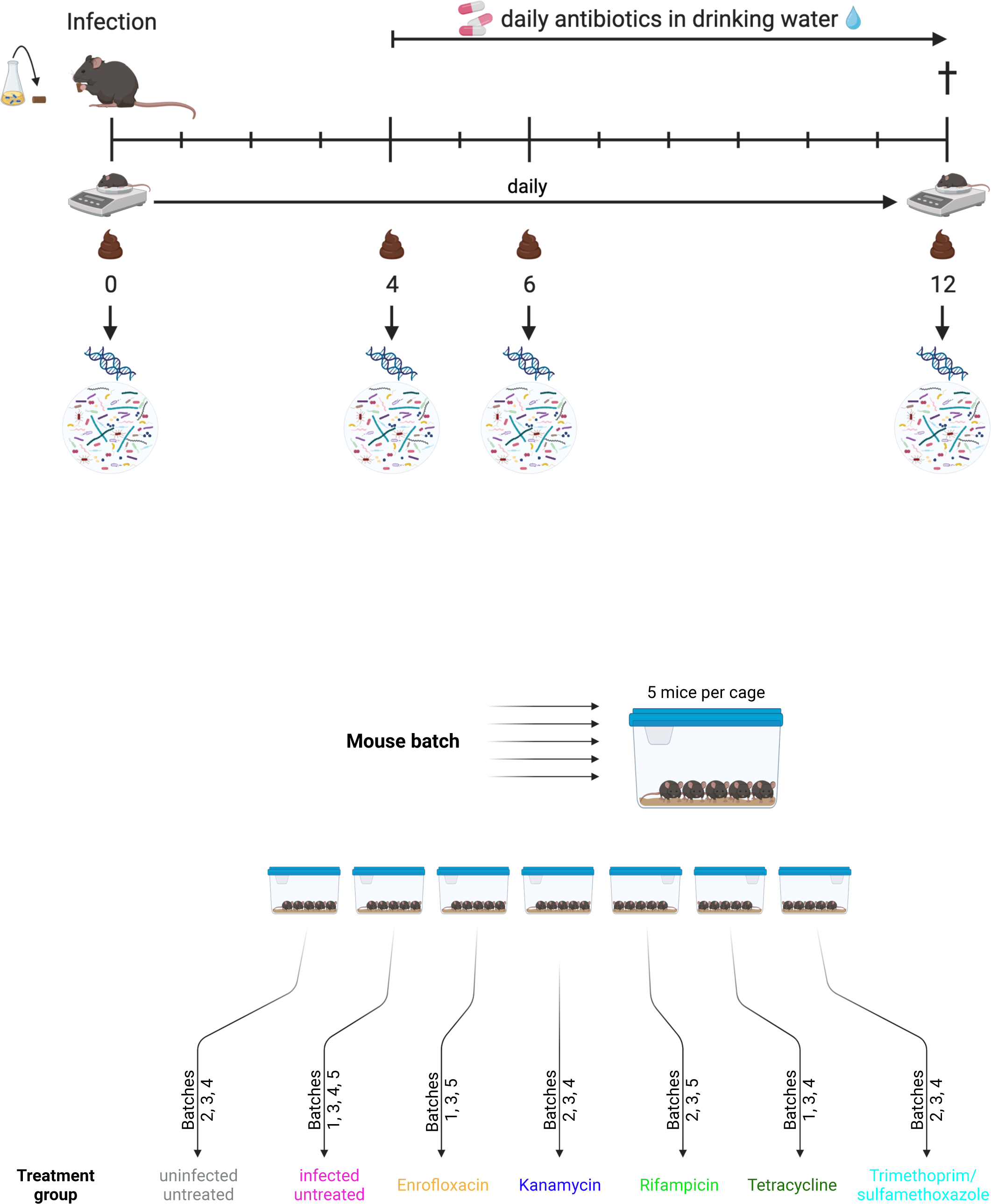
Experimental set-up. For each experiment, purchased mice were separated into groups of 5 mice per cage. Mice were infected with *C. rodentium* ϕ*stx2_dact_* by feeding on day 0 (D0), and weight was monitored daily (16). A group of control mice was kept uninfected. Prior to infection and on days 4, 6, and 12 (D4, D6, D12) post-infection, fecal samples were collected from which genomic DNA was isolated. From these samples, the V1/V2 regions of the 16S rDNA were amplified and sequenced for microbiota analyses. This figure was created with Biorender.com.

### Elimination of batch differences by mixing treatment groups

Differences in the intestinal microbiota composition of mice have been described and may depend on age, gender, genetic background, housing conditions, and others (30). We performed five experiments using independently purchased mouse batches (Batch 1-5). Each experiment consisted of different treatment groups to investigate the effect of *C. rodentium* ϕ*stx2_dact_* infection and antibiotics on the gut microbiota independently of the mouse batches (**Fig. 1**). We first assessed whether the mouse batches, although purchased from the same vendor and barrier, differed. For this purpose, we compared the fecal microbiota of all mice on Day 0 (D0). Permutational multivariate analysis of variance analysis (PERMANOVA) revealed that there were significant differences (p<0.001) between the different mouse batches at all taxonomic levels from sequence type up to phylum (**Table S2**). These differences are also represented in the non-metric multidimensional scaling (nMDS) plot (**Fig. S1A**). PERMANOVA comparisons between the different experiments showed that all batches were significantly different at the sequence type level (**Table S2**). Additionally, there were significant differences in sequence type richness (ST), evenness (J), and diversity **Fig. S1B**).

As every batch of mice was separated into groups of 5 mice, which were then infected and later divided into different treatment groups (by cage; see **Fig. 1**), we then determined whether the observed differences were also significant when the treatment groups were compared or whether the fact that the batches were all separated into the different treatment groups, was enough to eliminate the observed variability. Here, PERMANOVA revealed no differences in the microbiota composition at any taxonomic level (sequence type to phylum; **Table S2**). This can also be seen in the nMDS plot (**Fig. 2A**). Also, there were no significant differences in sequence type richness, evenness, or diversity (**Fig. 2B**). Hence, although there are significant differences in the purchased mouse batches before infection, splitting the mice into different treatment groups abrogated these variations.

**Fig. 2.**
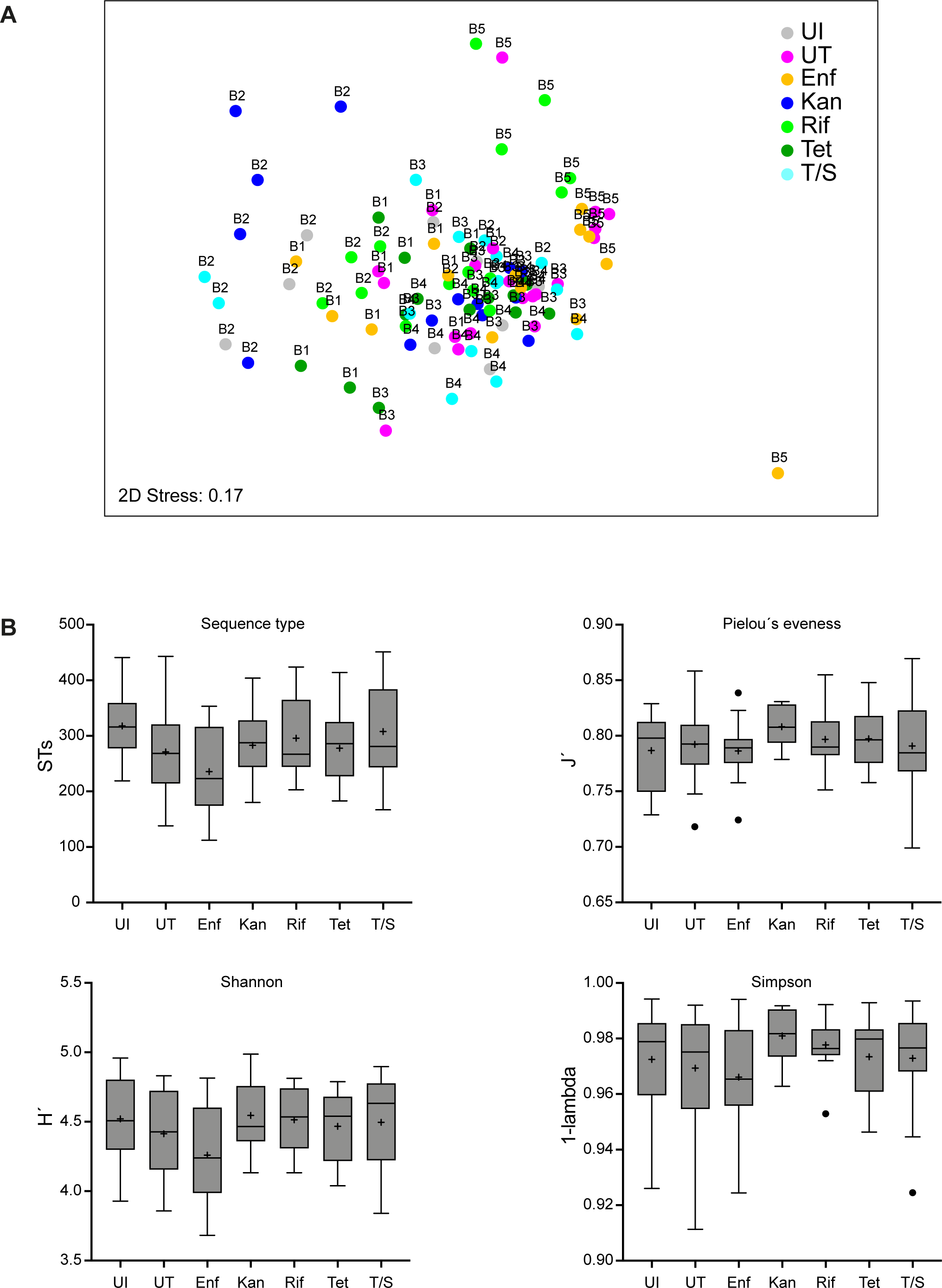
Batch mixing eliminates differences in treatment groups. **(A)** The global bacterial community structure in mice feces at the start of the experiment was assessed by non-metric multidimensional scaling (nMDS). The global community structure is based on standardized sequence type abundance data, and similarities were calculated using the Bray-Curtis similarity algorithm. Each mouse belonged to one of five batches (Batch) and was assigned to one treatment group (indicated by different colors). **(B)** The diversity of the different treatment groups is indicated by total sequence type number, Pielou’s evenness (J’), Shannon diversity (H’), and Simpsons diversity (1-λ), respectively, and was analyzed using sequence type relative abundance data as input. Data are based on an ordinary ANOVA analysis using Tukey’s test for multiple comparisons. The mean is indicated by + and the median by a black line. The box represents the interquartile range. The whiskers extend to the upper adjacent value (largest value = 75th percentile + 1.5 x IQR) and the lower adjacent value (lowest value = 25th percentile - 1.5 x IQR), and the dots represent outliers. There was no statistically significant difference in any of the tested indices between any treatment group.

### Influence of *C. rodentium* ϕ*stx2_dact_* infection on the fecal microbiota

To assess the impact of *C. rodentium* ϕ*stx2_dact_* infection on the fecal microbiota composition over time, samples taken from uninfected and infected untreated mice were compared. Uninfected untreated mice showed no significant changes in the microbiota composition down to the sequence type level over time (**Table S3**), which is reflected in the clustering of the samples in the nMDS plot (**Fig. S2A**). Some genera trended to be differentially distributed, but none were significantly different when corrected for multiple comparisons (**Fig. 3, Table S4**). Furthermore, there was no change in α-diversity at the sequence type level over time (sequence type richness (ST), Pielou’s evenness (J), diversity (Simpson index, 1-λ, and Shannon index H); **Fig. S2B).**

**Fig. 3.**
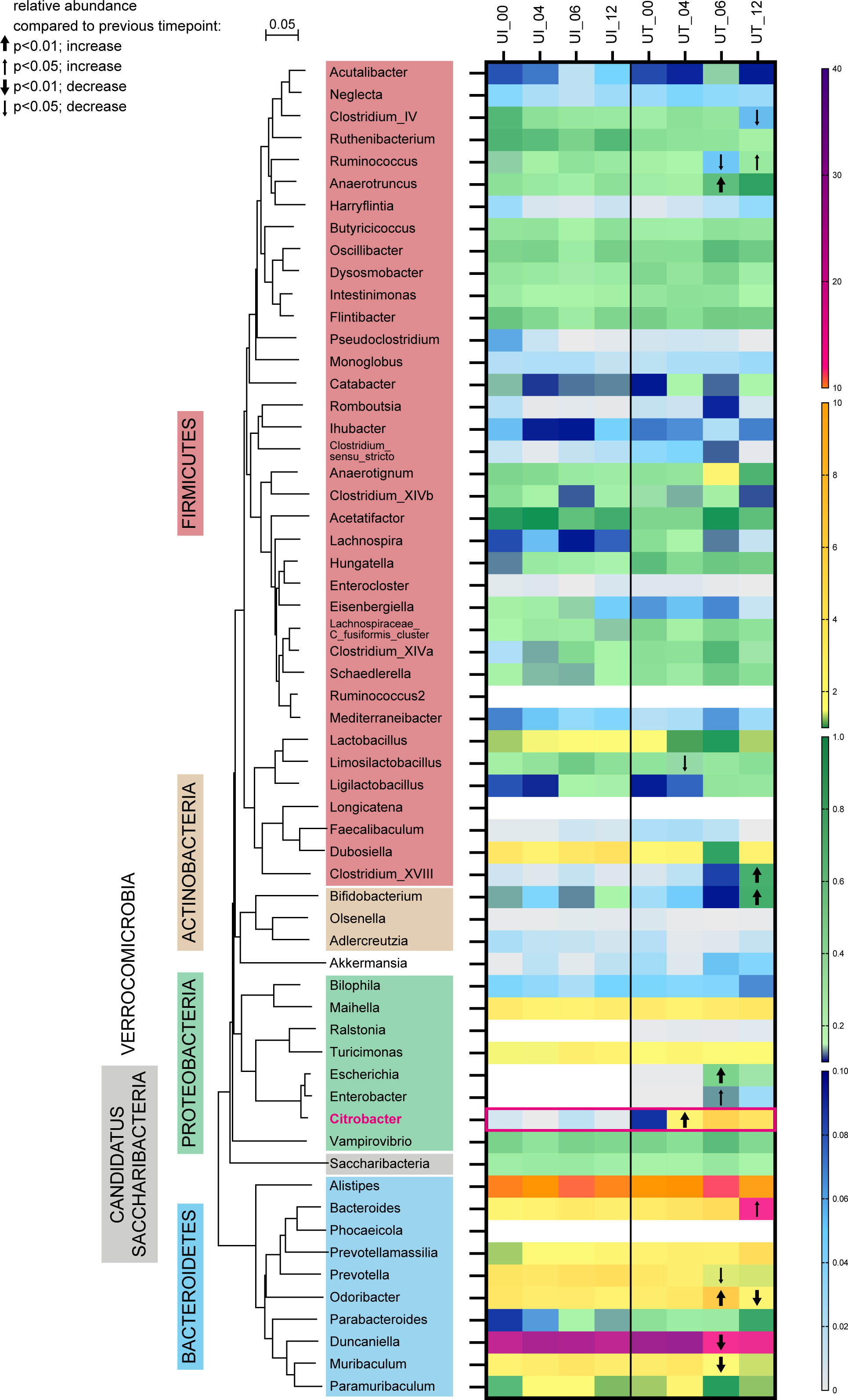
Heat map showing genera influenced by infection. The evolutionary history to the left was inferred using the Neighbor-Joining method and is based on representative nearly full-length 16S rRNA gene sequences for all genera given **(Table S5)**. The evolutionary distances were computed using the p-distance method and are given in units of the number of base differences per site. All ambiguous positions were removed for each sequence pair (pairwise deletion option). Evolutionary analyses were conducted in MEGA 7. The different phyla observed are indicated by color code. Only genera present in >10% of samples are further analyzed. The scale of the heatmap is indicated to the right and covers four orders of magnitude of mean relative abundance data. Changes over time were assessed by Kruskal-Wallis test with Benjamini-Hochberg corrections for multiple comparisons. Groups of samples were considered significantly different if the adjusted p-value was <0.05. Taxa differentially distributed over time were further assessed by Dunn’s post-hoc test. A significant change in abundance compared to the previous time point is indicated by a bold arrow if p<0.01 and a thin arrow if p<0.05. The arrow direction indicates an increase or decrease in abundance.

PERMANOVA analysis revealed significant community structure changes after infection with *C. rodentium* ϕ*stx2_dact_* from sequence type to the family level throughout the experiment and specific, tremendous changes upon infection from day 4 to day 6 visible at all taxonomic levels (**Table S3**). Concomitant with community structure changes, a clear increase in multivariate dispersion was observed (from 0.861 and 0.908 on day 0 (D0) and 4 (D4) respectively to 1.459 and 1.397 on day 6 (D6) and 12 (D12), respectively, **Fig. S3**). Only slight changes in diversity were observed and a significant increase in evenness (J) and diversity (1-λ) was visible only from D4 to D6 (**Fig. S2B**). The time course of *Citrobacter* abundance could be followed, which increased from a relative abundance of 2.0% on D4 to 5.0% on day 6 before slightly declining to 3.5% on D12 (**Fig. 3**). In addition to the significant change in *Citrobacter* abundance, significant changes in the abundance of 17 out of 74 genera or genus-level taxa (23%) were detected (**Fig. 3, Table S4**). The most prominent changes were observed for *Duncaniella* and *Muribaculum* of the Bacteroidetes six days post-infection (D6), where the relative abundance dropped to roughly 50%, whereas a further decrease during infection was not apparent. Similarly, *Prevotella* decreased significantly in abundance during early infection (D6) but not during late infection (D12). These observations contrast with *Odoribacter*, which increased significantly in abundance during early infection (D6), and *Bacteroides*, which increased significantly only in the late infection phase (D12). The increase in the abundance of Bacteroides could be further defined to the species level, where out of three species observed in the majority of samples, only *B. uniformis* increased significantly (see **Fig S4**). Outside the Bacteroidetes phylum, the effect on bacterial genera was minor (**Fig. 3, Table S4**). Of note, Enterobacteriaceae, related to *E. coli* but distinct from *Citrobacter,* increased from a mean relative abundance of <0.01% on D4 to 0.58% on D6. Together, these findings show that the *C. rodentium* ϕ*stx2_dact_* infection causes significant changes in the community structure of the murine intestinal flora.

### Consequences of antibiotic treatment on microbiota composition

We then investigated variations in the relative abundance of bacteria as a consequence of treatment with different antibiotics on D6 (short-term antibiotic treatment) and D12 (long-term antibiotic treatment) post-infection.

**Enrofloxacin** treatment was previously shown to clear *C. rodentium* ϕ*stx2_dact_* infection within two days. Unfortunately, while resolving the infection, the antibiotic-induced Shiga toxin expression and release resulted in severe kidney pathology, weight loss, and death (16). As expected, the abundance of *Citrobacter* was greatly reduced after treatment onset (from a mean of 3.2% on D4 to 0.01% on D6), and the pathogen was eliminated afterwards (**Fig. 3, Table S4**). The reduction of *Citrobacter* abundance was accompanied by a significant change in community structure as evidenced by PERMANOVA analysis (**Table S3**; see also visualization in the nMDS plot **Fig. S5A**), with a slight decrease in evenness from D6 to D12 (0.815 ± 0.056 on D6 to 0.768 ± 0.062 on D12), but no significant effect on richness and diversity (**Fig. 4A**). The multivariate dispersion increased tremendously from 0.986 and 1.063 on D0 and D4 respectively to 1.73 and 1.818 on D6 and D12, respectively (**Fig. S3**), the highest heterogeneity observed here. Fifty-two of 71 genera (73%) showed a significant change in their abundance over time (**Table S4**). Forty-eight of these were affected during short treatment (D4 vs. D6), but only 5 during long-term treatment (D6 vs. D12) with enrofloxacin. Interestingly, the majority of genera of the Bacteroidetes, Proteobacteria, and Actinobacteria phyla decreased considerably in abundance during early treatment (**Fig. 6**), with all three *Bacteroides* species were practically eliminated already on D6 (**Fig. S4**). In contrast, no clear trend was observed within the Firmicutes (**Fig. 6**). Several Ruminococcaceae (e.g., *Harryflintia, Anaerotruncus*) as well as *Oscillibacter* and *Dysosmobacter* increased in relative abundance after short-term antibiotics treatment (D6), but declined later on, whereas an increase in relative abundance of Clostridiales was observed (**Fig. 6, Table S4**). This trend was also visible in the genera Lachnospiraceae and specifically in unclassified Lachnospiraceae. They comprised a mean of 13.8% on D4 before enrofloxacin treatment and increased significantly to a mean of 48.9%. These differences were also visible when higher taxa (families to phylum) were analyzed (**Fig. S6**).

**Fig. 4.**
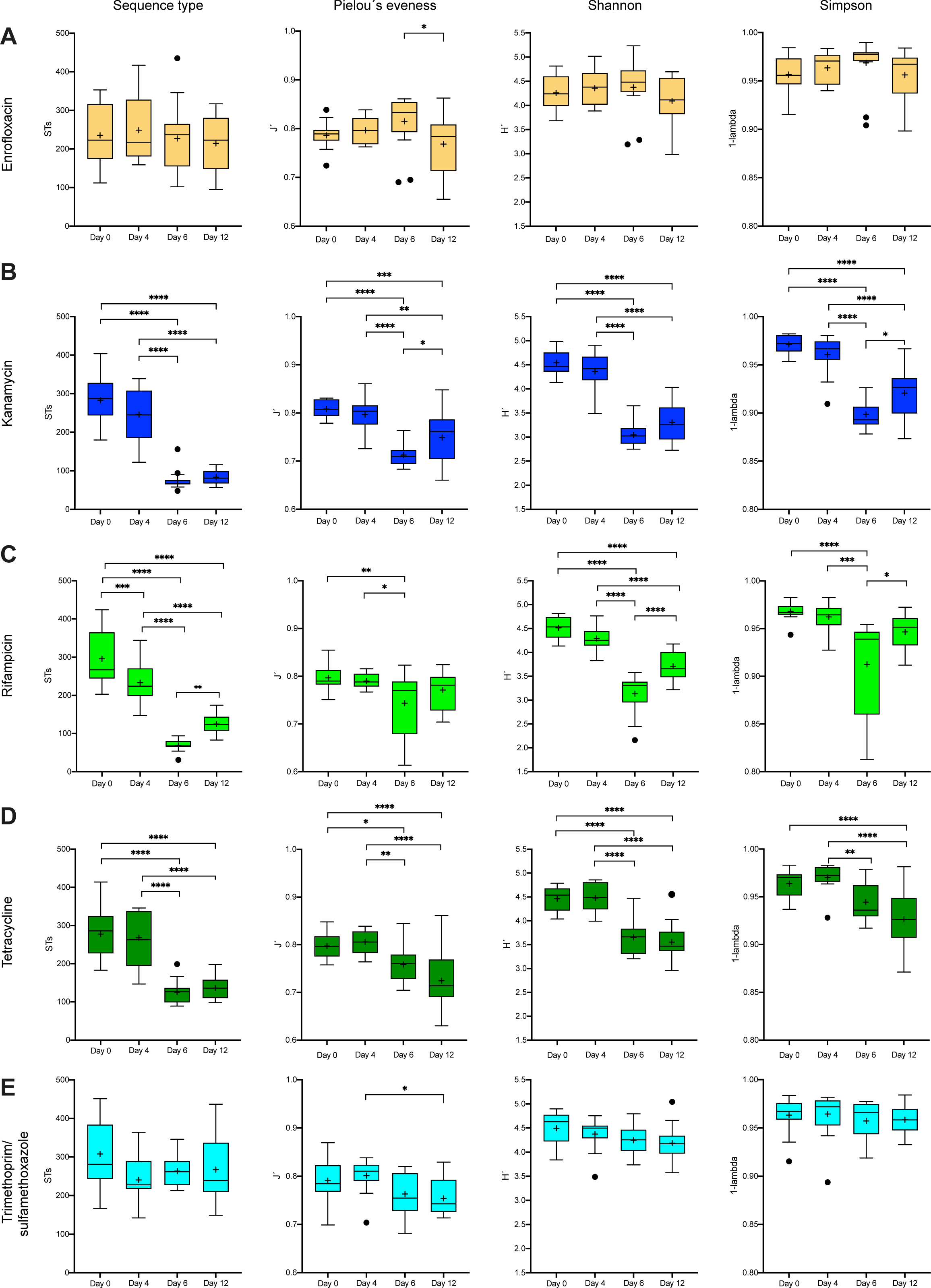
Bacterial community diversity dependent on antibiotic treatment. Diversity is indicated by total sequence type number, Shannon diversity (H’), Simpson’s diversity (1-λ), and Pielou’s evenness (J’), respectively, and was analyzed using sequence type relative abundance data as input. Differences in diversity were analyzed using a mixed effects model and multiple comparisons were corrected using the Tukey test (A, enrofloxacin; B, kanamycin; C, rifampicin; D, tetracycline; E, trimethoprim/sulfamethoxazole) separately over time. Statistically significant differences are indicated as *p<0.05, **p<0.01, ***p<0.001 or ****p<0.0001. The mean is indicated by + and the median by a black line. The box represents the interquartile range. The whiskers extend to the upper adjacent value (largest value = 75th percentile + 1.5 x IQR) and the lower adjacent value (lowest value = 25th percentile - 1.5 x IQR) and the dots represent outliers.

Treatment with **kanamycin** also allowed complete elimination of *C. rodentium* ϕ*stx2_dact_* on day 6 post-infection with low colon pathology and no kidney damage (16). However, kanamycin caused the most dramatic changes in microbiota composition (**Table S3; Fig. S5B**), with 60 out of 65 genera (92%) showing a significant change (mainly a reduction) in relative abundance over time (**Fig. 6, Fig. S6, Table S4**). In contrast to enrofloxacin, which had a minor impact on diversity and induced only a slight decrease in evenness, kanamycin significantly influenced taxon richness (from 246±69 on D4 to 75±25 on D6), evenness (from 0.797±0.035 to 0.713±0.025), and diversity (H: from 4.36±0.39 to 3.05±0.23; 1-λ: from 0.961±0.020 to 0.899±0.015) (**Fig. 4B**). A slight recovery in evenness and diversity was observed during long-term kanamycin treatment (D12) (**Fig. 4B**). Also, in contrast to enrofloxacin a decrease rather than an increase in dispersion was observed (**Fig. S3**). In accordance with our previous data (16) and as observed for enrofloxacin, the abundance of *Citrobacter* was tremendously reduced after short-term kanamycin treatment and completely abolished on D12 (**Fig. 6, Fig. S7**). Furthermore, all other proteobacterial genera significantly diminished during short-term antibiotic treatment, and bacterial reads that could be classified to any proteobacterial class were absent on D12. Similarly, both actinobacterial genera (*Adlercreutzia* and *Bifidobacterium*) as well as unclassified *Eggerthellaceae* were nearly abolished on D6 with only a few reads remaining in some communities (**Fig. 6, Table S4**). Nearly all Clostridiales genera were also practically eliminated already on D6, and unclassified Ruminococcaceae, unclassified Lachnospiraceae, or unclassified Clostridiales followed the same trend with a significant reduction during short-term antibiotic treatment. An exception were bacteria with similarity in sequence to *Clostridium fusiformis* of the Lachnospiraceae, which increased in relative abundance and reached a mean of 1.4%. The Erysipelotrichiaceae genera *Faecalibaculum* and *Duboisiella* also significantly increased in relative abundance upon kanamycin treatment (**Fig. 6, Fig. S7, Table S4**). Members of the Bacteroidetes showed a mixed behavior, where specifically *Bacteroides* increased by more than one order of magnitude in relative abundance during short-term antibiotic treatment (from 2.5 to 31.9%). The most extreme change was observed for *B. acidifaciens,* which increased from a mean of 0.5% on D4 to a mean of 24.4% on D6, whereas Bacteroides 11 was only slightly affected (**Fig. S4**). Also, *Parabacteroides* increased from below 0.1% before antibiotic treatment to a mean of 5.2% relative abundance on D6 and 7.8% on D12, whereas e.g., *Odoribacter* and *Muribaculum* decreased (**Fig. 6, Table S4**). These differences in abundance upon treatment were also observed at higher taxonomic levels where overall only Bacteroidales showed an increase in relative abundance, whereas the abundance of all other phyla decreased (**Fig. S6**).

**Rifampicin** also allowed survival and prevented kidney damage by the infection with *C. rodentium* ϕ*stx2_dact_*, but overall, the colon pathology was somewhat higher, as the pathogen was not fully eliminated (16). Microbiota analysis confirmed this result, and *Citrobacter* remained as an important member of the microbial community, although at a relatively low abundance of 0.2% after 6 and even 12 days (**Fig. 6**). Rifampicin treatment was accompanied by a substantial reduction in richness (from 233±52 on day 4 to 69±15 on D6) and diversity (H: from 4.291±0.246 to 3.134±0.408; 1-λ: from 0.962±0.014 to 0.913±0.054) that recovered slightly but significantly over treatment time (richness: 125±27 on D12; H: 3.710±0.287; 1-λ: 0.947±0.020) (**Fig. 4C**). There was a slight increase in multivariate dispersion concomitant with antibiotic treatment (**Fig. S3**). PERMANOVA shows that rifampicin treatment resulted in significant changes in microbiota composition over time (**Table S3**), and the nMDS plot revealed a slight recovery of microbiota composition on D12 (**Fig. S5C**). This was reflected in the number of genera differentially distributed where out of 60 genera, 47 (83%) were affected during early antibiotic treatment (D4 to D6) and 25 during late antibiotic treatment (D6 to D12) (**Fig. 6, Table S4**). As observed for kanamycin treatment, most Clostridiales genera clearly diminished in relative abundance. Also, unclassified Lachnospiraceae diminished dramatically in relative abundance by two orders from D4 to D6 (**Table S4**). However, they recovered to pre-treatment levels on D12. Interestingly, recovery was also observed for a variety of Ruminococcaceae and Lachnospiraceae genera (see **Fig. 6, Fig. S7**). In contrast, Erysipelotrichiaceae (*Faecalibaculum* and *Clostridium* XVIII) increased significantly after rifampicin treatment. Such an increase in relative abundance was also evident for various proteobacterial genera. However, during extended treatment, they regained their original relative abundance levels. Enterobacteriaceae related to *E. coli* and distinct from *Citrobacter* increased from a mean relative abundance of <0.01% on D4 to 9.2% on D6 and then dropped to 0.32% on D12. The effect of rifampicin on Bacteroidetes was typically negative and resulted in relative depletion of *Alistipes*, *Prevotella*, *Odoribacter*, and *Parabacteroides,* usually by at least one order of magnitude (**Fig. 6, Fig. S7, Table S4**). Accordingly, depletion was observed for the whole Bacteroidales class as well as the Clostridiales, whereas Erysipelotrichiales followed opposing abundance effects (**Fig. S6**).

Similar to kanamycin, **tetracycline** was also able to fully eradicate *Citrobacter* colonization on D6, allowed murine survival, and abolished colon and kidney damage (16). The depletion could be confirmed here by microbiota analysis, where the relative abundance of *Citrobacter* was 0.004% on D6, with no *Citrobacter* detectable on D12 (**Fig. 6, Table S4**). Diversity changes during tetracycline treatment were prominent with a severe decline in richness (268±72 to 125±31), evenness (J: 0.806±0.023 to 0.758±0.039), and diversity (H: 4.479±0.299 to 3.652±0.362; 1-λ: 0.970±0.014 to 0.945±0.019) from D4 to D6 (**Fig. 4D**). A clear shift in microbiota composition was observable and while samples obtained on D0 and D4 clustered together in the nMDS plot, the samples for both D6 and D12 show distinct, separate clustering (**Fig. S5D**), suggesting successive shifts in microbiota composition throughout treatment. These changes were statistically significant, as evidenced by PERMANOVA (**Table S3**). Interestingly, tetracycline addition did not affect multivariate dispersion (**Fig. S3**). A total of 52 of 66 genera (79%) were influenced by tetracycline treatment, with 47 being impacted from D4 to D6 and 12 genera from D6 to D12 (**Fig. 6, Fig. S7, Table S4**). The application of tetracycline resulted in a rapid depletion and elimination of Lactobacillaceae, Actinobacteria, and Proteobacteria (**Fig. 6, Fig. S6, Table S4**). Also, most Clostridiales genera were negatively affected. However, *Ruthenibacterium* and Lachnospiraceae of the *C. fusiformis* cluster showed an increase during early treatment. Bacteroidetes showed different oscillating behavior. All three Muribaculaceae genera (*Duncaniella*, *Muribaculum,* and *Paramuribaculum*) decreased under tetracycline treatment, with both *Muribaculum* and *Paramuribaculum* recovering during extended treatment. Similarly, the relative abundance of *Prevotellamassilia*, *Prevotella,* and *Odoribacter* decreased, and that of *Prevotellamassilia* and *Prevotella* returned to higher relative abundance levels during extended treatment. In contrast, *Alistipes* and *Bacteroides* showed an extreme initial relative abundance increase during tetracycline treatment (**Fig. 6, Fig. S7, Table S4**). A detailed analysis of the species level revealed that only *Bacteroides* 11 and *B. acidifaciens* contributed to this overall increase of the genus, with *B. acidifaciens* increasing from 0.6% relative abundance on D4 to 40.2% on D12. This contrasts with the severe depletion observed for *B. uniformis* **(Fig. S4)**.

In the case of **trimethoprim/sulfamethoxazole,** complete elimination of *C. rodentium* ϕ*stx2_dact_* was only observed on D12 but not on D6 post-infection, and this was sufficient to reduce but not abolish colon pathology or kidney damage (16). Accordingly, microbiota analysis revealed significant *Citrobacter* levels on D6 (0.014%) (**Fig. 6, Fig. S7, Table S4**). Changes in diversity and richness were minor, and only slight changes in evenness were recorded as significant (**Fig. 4E**). Dispersion increased after antibiotic treatment (from 0.946 and 0.943 on D0 and D4, respectively, to 1.579 and 1.612 on D6 and D12, respectively, **Fig. S3**). Also, the fecal microbiota composition changed significantly (**Fig. 6, Fig. S7, Table S3**) but less tremendously compared to, for example, kanamycin and tetracycline. Only 31 of 69 genera (45%) were significantly affected in their abundance. Of these, 21 were affected during early treatment and only five during extended treatment. For trimethoprim/sulfamethoxazole, the most prominent effect observed was a decrease in the relative abundance of various Bacteroidetes genera such as *Prevotella*, *Odoribacter*, *Duncaniella*, *Muribaculum,* and *Paramuribaculum*. Clostridiales genera were only slightly affected, whereas Lactobacillaceae increased in relative abundance (**Fig. 6, Fig. S7, Table S4**).

In summary, both short and long-term antibiotic treatment resulted in significant and global shifts in microbiota composition (**Fig. 6, Fig. S7**, **Table S4**), which were much more dramatic compared to those observed during infection (**Fig. 3, Fig. S7**). Furthermore, these changes were highly specific for each tested antibiotic, with kanamycin having the most prominent effect. In contrast, trimethoprim/sulfamethoxazole triggered relatively minor differences, and no substantial overlaps between treatment groups were observed (**Fig. 5**, **Fig. 6**).

**Fig. 5.**
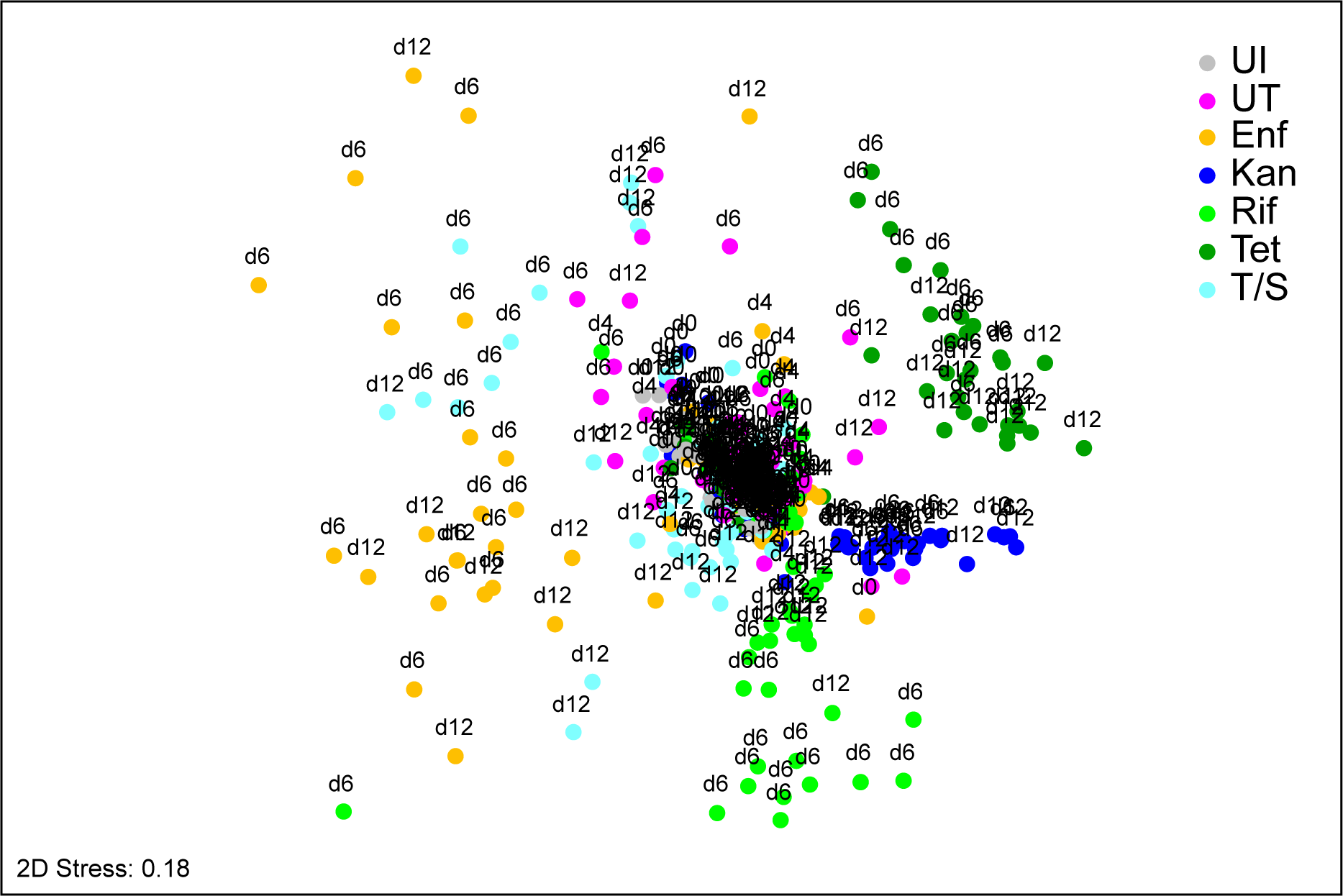
Differences in global bacterial community structure in mice feces upon infection and subsequent antibiotic treatment. The global bacterial community structure was assessed by non-metric multidimensional scaling (nMDS) and is based on standardized sequence type abundance data. Similarities were calculated using the Bray-Curtis similarity algorithm. All treatment groups except control mice (UI) were infected with *C. rodentium* ϕ*stx2_dact_* on day 0. Treatment groups that received antibiotics from day 4 post-infection are indicated by Enf (enrofloxacin), Kan (kanamycin), Rif (rifampicin), Tet (tetracycline), or T/S (trimethoprim/sulfamethoxazole). Treatment group ‘UT’ remained untreated. The labels indicate the day post-infection.

**Fig. 6.**
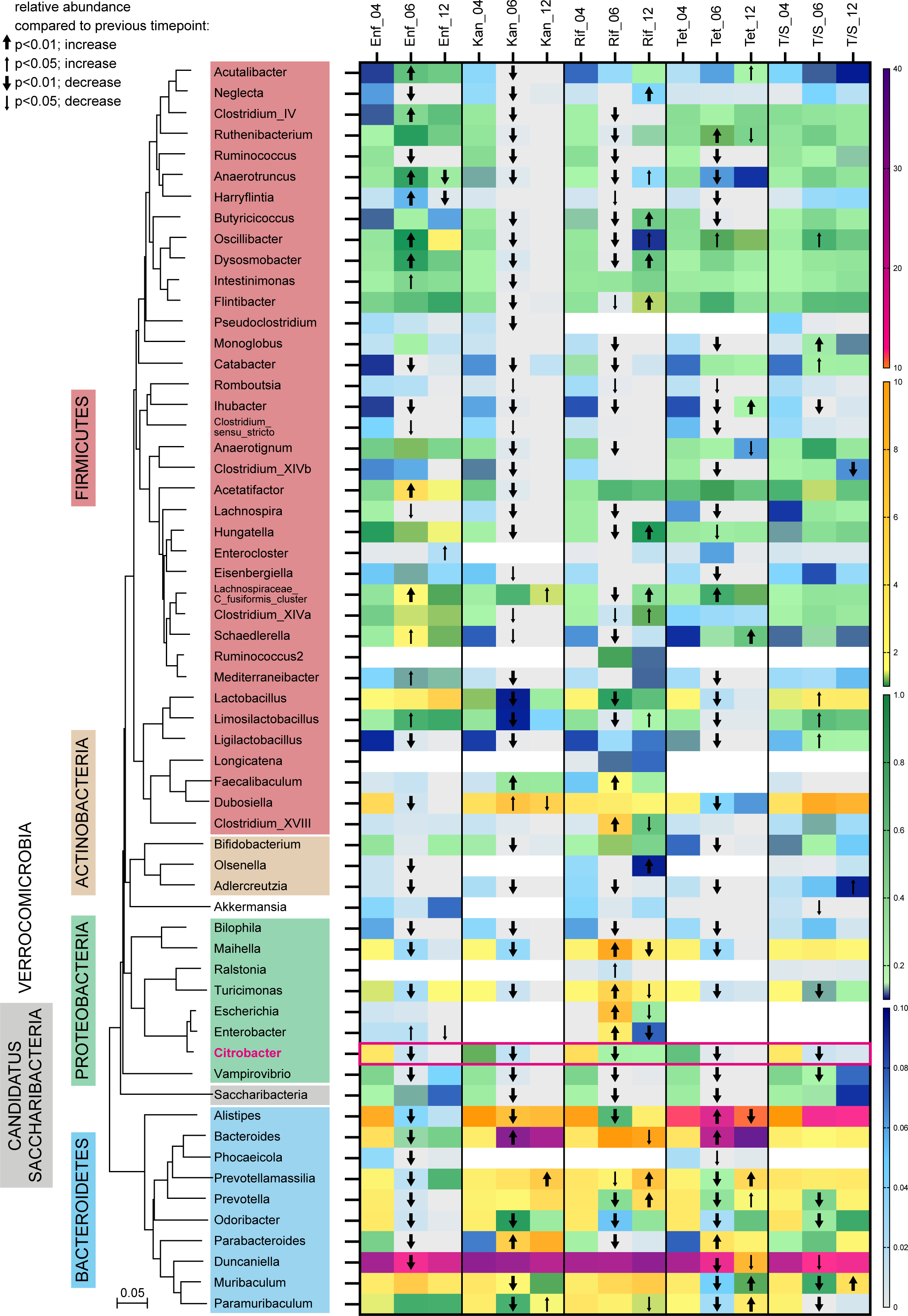
Heat map showing genera influenced by antibiotic treatment post-infection. The evolutionary history to the left was inferred using the Neighbor-Joining method and is based on representative nearly full-length 16S rRNA gene sequences of representatives for all genera given **(Table S5)**. The evolutionary distances were computed using the p-distance method and are in the units of the number of base differences per site. All ambiguous positions were removed for each sequence pair (pairwise deletion option). Evolutionary analyses were conducted in MEGA 7. The different phyla observed are indicated by color code. Only genera present in >10% of a given treatment group are further analyzed. The scale of the heatmap is indicated to the right and covers four orders of magnitude of mean relative abundance data. Changes over time were assessed by Kruskal-Wallis test with Benjamini-Hochberg corrections for multiple comparisons. Groups of samples were considered significantly different if the adjusted p-value was <0.05. Taxa differentially distributed over time were further assessed by Dunn’s post-hoc test. A significant change in abundance compared to the previously indicated time point is indicated by a bold arrow if p<0.01 and a thin arrow if p<0.05. The arrow direction indicates an increase or decrease in abundance.

## DISCUSSION

Several recent studies described the impact of single and, in some cases, combinations of antibiotics on the intestinal microbiota in mice (30, 32–36, 39). However, a detailed comparative analysis addressing the impact of different classes of antibiotics on the gut microbiota during treatments to eliminate enteric bacterial pathogens has not been performed. Here, we describe that infection with *C. rodentium* ϕ*stx2_dact_* used to mimic EHEC infections in mice, causes significant shifts in the relative abundance of members of the fecal microbiota. However, the overall effect of the infection alone was minor compared to that triggered by the treatment with antibiotics.

Mice in which the infection remained untreated displayed a significant increase of *Citrobacter* with a maximum relative abundance of ∼5% on day 6. This corresponds to the relative abundance previously reported for the intestinal lumen (40). However, the mucosal relative abundance appears to reach higher values (38, 41). While no alterations in microbial diversity upon *C. rodentium* infection were reported in the literature (40), minor variations in microbiota composition were detected (38, 40–43). Lupp *et al.* indicated a bloom of Enterobacteriaceae upon infection, yet Hoffmann *et al.* suggested that this bloom at the mucosa was mainly due to an expansion of *Citrobacter* itself rather than of other family members (41, 42). Also, Hopkins *et al.* reported an increase in Enterobacteriaceae, which was restricted to the mucosa, however, without analyzing whether this bloom was solely due to *Citrobacter* (38). We could clearly show that on day 6 post-infection, an increase in Enterobacteriaceae distinct to *Citrobacter* occurred, probably due to *Citrobacter* creating a niche for those Enterobacteriaceae.

The most prominent effect of infection observed in our study was a decrease in Bacteroidetes, specifically due to a decrease in relative abundance of Muribaculaceae (**Fig. S7**), a bacterial family just recently defined and before subsumed into the Porphyromonadaceae (44). One previous report also described a decrease in Porphyromonadaceae and Prevotella, another a decrease in *Lactobacillus* abundance upon murine infection with wildtype *Citrobacter* (42, 43) which is similar to the Stx-expressing *Citrobacter* variant used here. However, a detailed comparison with other reports is difficult as they limited the analysis to higher taxonomic levels (40), analyzed very few animals (38), or the microbiota of the uninfected, naïve mice used in the different studies varied significantly, which is likely to affect the pathogen colonization and pathogen-triggered changes of the microbiota (35). For example, Hoffmann *et al.* reported a significant increase in Deferribacteriaceae (42), which were absent from the microbiota of mice analyzed here.

In fact, the naïve microbiota of mice used here varied significantly depending on the mouse batch, even when obtained from the same supplier. While the significance of these community differences was eliminated by mixing different mouse batches to create treatment groups, differences remained at the individual mouse level and may create disparities in colonization resistance mediated by the naïve microbiota (43). As example, resistance to *C. rodentium* and EHEC infection has previously been associated with higher diversity and abundance of butyrate-producing bacteria (29, 45) and higher concentrations of SCFAs (29), and administration of butyrate reduced pathogen-mediated intestinal damage (46). Also, enhancement or erosion of the mucus layer by commensals may contribute to the colonization capability (47). Moreover, EHEC and *Citrobacter* virulence gene expression is controlled by microbiota-derived substances, which will also influence colonization and associated microbiota alterations (47, 48).

Infections with EHEC are commonly not treated with antibiotics due to the fear of antibiotic-induced Shiga toxin production (7–9). However, as the *E. coli* O104:H4 outbreak in Northern Germany in 2011 has shown, the necessity may arise to treat patients with antibiotics to prevent deadly outcomes. The effects of antibiotics from different classes on the induction of *stx* expression, however, vary greatly. Antibiotics that interfere with DNA replication (e.g., enrofloxacin, ciprofloxacin) induce the bacterial SOS response and tend to induce Shiga toxin synthesis *in vitro*. In contrast, antibiotics inhibiting protein synthesis (e.g., kanamycin, tetracycline) or blocking bacterial transcription (e.g., rifampicin) consistently showed no *stx* induction (7, 9, 13, 16, 49) and enhanced survival of EHEC- or *C. rodentium* ϕ*stx2_dact_*-infected mice (16, 50). In particular, tetracycline and kanamycin cleared an infection with *C. rodentium* ϕ*stx2_dact_* without causing kidney damage (16) whereas rifampicin only reduced the *Citrobacter* load, but limited kidney damage (16). Importantly, several patients who developed HUS during the *E. coli* O104:H4 outbreak were treated with rifaximin, a rifampicin derivative. All patients survived and had fewer seizures than those not treated (51), suggesting that these antibiotics may provide promising options for life-threatening EHEC infections.

The use of antibiotics is well-known to cause dysbiosis (52, 53). However, little information was available regarding the influence of individual antibiotics on the fecal microbial community structure in a comparable EHEC infection setting. Here, we showed that mice treated with individual antibiotics of different classes showed large, antibiotic-specific shifts in microbiota composition and varied in their response to long-term antibiotic treatment. We showed here that the administration of kanamycin, tetracycline or rifampicin as promising treatment options all resulted in significant abundance changes of at least 75% of genera and genus-level taxa observed and caused a significant reduction in diversity. This dysbiosis may trigger adverse effects, including the opening up of niches for infection with or outgrowth of pathobionts such as *Clostridioides difficile* and *Enterococcus faecalis* (5, 6), which should be considered for clinical applications.

Several recent studies have also investigated the effects of single antibiotics or antibiotic cocktails on the intestinal microbiota of naïve mice. However, some of these studies used very few animals in the different treatment groups (31, 32, 34), such that the significance of identified differences can only poorly be assessed. For example, Sun *et al.* (31) used only five mice per treatment group and detected no changes in microbiota composition upon enrofloxacin treatment at the phylum level but an increase in Prevotellaceae and Rikenellaceae and decrease in Bacteroidaceae families (31). This contrasts with our report, where a significant impact on nearly all Bacteroidetes genera and all Bacteroidetes families and various other genera could be evidenced with 3-fold that sample size. Another study used qPCR on a restricted number of taxa to survey the microbiota. However, the results (54) do not correspond to those of other studies, possibly because the method does not reach the accuracy of 16S rDNA amplicon sequencing or metagenomic analyses. In a metagenomic study assessing the effects of tetracycline, Yin *et al.* observed a significant decrease in the abundance of Firmicutes together with an increase in Bacteroidetes, in accordance with our results (33), but Zhao *et al.* obtained slightly different results in their study, which, however, was carried out with an extremely small sample size (34). Namasivayam *et al.* evaluated the effect of anti-tuberculosis therapy but also of rifampicin alone in small treatment groups (32). Most of the observed community changes were due to rifampicin, and *Alistipes*, *Erysipelotrichiaceae,* and *Parabacteroides* trended to be depleted. Mullineaux-Sanders *et al.* (47) used six animals per treatment group and showed that kanamycin treatment exhibited severe effects and that the bacterial communities were highly dominated by Bacteroidetes genera, similar to the situation observed in our study. Korte *et al.* also evaluated trimethoprim/sulfamethoxazole, which induced minimal changes in the community composition (35). Here, we also showed that trimethoprim/sulfamethoxazole did not affect the diversity and caused milder alterations of the microbiota overall. However, this antibiotic was less effective, as it only slowly eliminated the pathogen and reduced but did not abolish Stx-mediated kidney damage (16). Overall, our data confirmed that trimethoprim/sulfamethoxazole and enrofloxacin have a less disruptive effect on the microbiota than tetracycline and kanamycin.

Other studies also included mice with a different, rather unusual naïve mouse microbiota, which also hampered a direct comparison of the antibiotic effects. Very recently, Grabowski *et al.* (30) aimed to analyze the effect of enrofloxacin, however, the control mice exhibited a tremendously high amount of Carnobacteriaceae (∼30%) and Pseudomonadaceae (10%) as unusual gut colonizers, which were rapidly depleted preventing any detailed evaluation of antibiotic effects. Severe and long-term changes in the intestinal microbiota were also observed with a combination of four different antibiotics (ampicillin, vancomycin, metronidazole, and neomycin) (30, 36, 37, 39). Here, an increase in *Enterococcus* and a decrease in probiotics-related genera such as *Lactobacillus* was reported as common across individual and mixed antibiotic treatments (37). However, *Enterococcus* is not a common member of the murine microbiome and was present here at very low abundance in only a small subset of mice.

The comparison with recent studies made evident that reported antibiotic-mediated effects on the murine microbiota differ significantly, depending not only on the antibiotic and the treatment conditions but considerably on the composition of the naïve microbiota, the sample size used for analysis, and the sensitivity of the applied microbiota analysis method. All these factors contribute to seemingly inconsistent results between studies. Moreover, additional information is required about inhibitory and enhancing effects associated with individual antibiotics, such as concentrations of metabolites controlling microbial growth and host immune responses, as well as the expression of virulence-relevant genes of intestinal pathogens. A more detailed understanding of the effects of antibiotic treatment on different members of the gastrointestinal microbiota will be of great importance to address these issues.

## MATERIAL AND METHODS

### Animal ethics

C57BL/6Rj mice were housed under pathogen-free conditions in accordance with FELASA recommendations in the BSL3 animal facility of the Helmholtz Centre for Infection Research, Braunschweig. The protocol was approved by the Niedersächsisches Landesamt für Verbraucherschutz und Lebensmittelsicherheit: permit no. 33.19-42502-04-16/2124. Animals were treated with appropriate care, and all efforts were made to minimize suffering. Food and water were available *ad libitum* throughout the experiment.

### Animal infections

Six-week-old female C57BL/6Rj mice purchased from Janvier Labs (Le Genest-Saint-Isle, France) barrier A02 were infected with 5 x 10^8^ CFU *C. rodentium* DBS770 (*C. rodentium* ϕ*stx2_dact_*) following the feeding protocol described in Flowers *et al.* (55) or left uninfected. From 4 days post-infection, drinking water was supplemented with 2% glucose and either of the following antibiotics: enrofloxacin (0.25 mg/ml), kanamycin (2.6 mg/ml), tetracycline (1 mg/ml), rifampicin (1 mg/ml), or trimethoprim/sulfamethoxazole (Trimetotat oral suspension 48% (Livisto)). Supplemented water was exchanged daily to ensure continuously high levels of antibiotics. Mice were weighed daily.

### Genomic DNA isolation from feces

For collection of fecal pellets from individual mice, animals were separated into boxes for up to 30 min. Samples for gDNA isolation were collected in Lysing matrix D tubes on D0 (before infection) and on D4, D6, and D12 post-infection. Genomic DNA was isolated using the FastDNA^TM^ Spin Kit and a FastPrep-24 bead beating grinder (MP Biomedicals, Germany) and eluted in 100 µl H_2_O_dd_. DNA concentrations were determined using a NanoDrop™ One/One^C^ spectrophotometer (ThermoFisher Scientific).

### 16S rDNA amplification and sequencing

A 2-step PCR-approach was used to amplify the V1-V2 variable region of the 16S rRNA gene. PCR with primers 27Fbif and 338R containing part of the sequencing primer sites as short overhangs (given in italics) (*ACGACGCTCTTCCGATCT*AGRGTTHGATYMTGGCTCAG and *GACGTGTGCTCTTCCGATCT*TGCTGCCTCCCGTAGGAGT, respectively) was used to enrich for target sequences (20 cycles). A second amplification step of 10 cycles added the two indices and Illumina adapters to amplicons (56). Amplified products were purified, normalized, and pooled using the SequalPrep Normalization Plate (ThermoFisher Scientific) and sequenced on an Illumina MiSeq (2X300 bases, San Diego, USA). Demultiplexed raw data for all the amplicon sequencing pair-end datasets are publicly available at the NCBI Sequence Reads Archive (SRA) under BioProject accession number PRJNA1011327.

### Bioinformatic and statistical analysis

The fastQ files were analyzed with the dada2 package version 1.12.1 in R (57). The quality-trimming and filtering steps were performed using the filterAndTrim function. Forward and reverse reads were trimmed on the 5’-end by 20 and 19 bases, respectively. Reads were truncated to a length of 240 bases and a maximum of 2 expected errors per read was permitted. After denoising and paired-end reads merging, chimeras were removed. Remaining non-bacterial sequences (eukaryota, mitochondria, chloroplast) were manually deleted. Overall, 7,654,419 bacterial 16S rDNA sequence counts were obtained with a mean of 18,669 ± 8,447 reads per sample (**Table S1**). All samples were re-sampled to equal the smallest library size of 6,708 reads using the phyloseq package returning 3,098 sequence types (58) (**Table S1**). Sequence types were annotated based on the naïve Bayesian classification with a pseudo-bootstrap threshold of 80% using RDP set18 (59) (**Table S1**). Sequence variants were then manually analyzed against the RDP database using the Seqmatch function to define the discriminatory power of each sequence type. Species level annotations were assigned to a sequence variant when only 16S rRNA gene fragments of previously described isolates of a single species were aligned with a maximum of two mismatches with this sequence variant (60). Relative abundances (in percentage) of sequence types, species, genera, families, orders, classes and phyla were used for downstream analyses. Calculation of diversity indices (species richness ST, Shannon diversity index H, Pielous evenness J, Simpson diversity index 1-λ) and multivariate analyses were performed using PRIMER (v.7.0.11, PRIMER-E, Plymouth Marine Laboratory, UK), whereas univariate analyses were performed using Prism 9 (Graphpad Software, Inc.).

Differences in diversity indices between the different mouse batches (**Fig. S1B**) and the different treatment groups (**Fig. 2B**) were tested for by ordinary ANOVA using the Holm-Sidak test for multiple comparisons. Changes in diversity over time were analyzed using a mixed effects model. Tukey’s test was used for multiple comparisons.

The data matrices comprising 3,098 sequence types, 126 genera, or other taxa were used to construct sample-similarity matrices applying the Bray-Curtis algorithm, where samples were ordinated using non-metric multidimensional scaling (nMDS) with 50 random restarts (61). Significant differences between *a priori* predefined groups of samples were evaluated using Permutational Multivariate Analysis of Variance (PERMANOVA), allowing for type III (partial) sums of squares, fixed effects sum to zero for mixed terms. Monte Carlo p-values were generated using unrestricted permutation of raw data (62). Groups of samples were considered significantly different if the p-value was <0.05. The abundances of taxa present in the community of at least 10% of the samples were compared by the Kruskal-Wallis test with Benjamini-Hochberg corrections for multiple comparisons (63). Groups of samples were considered significantly different if the adjusted p-value was <0.05. Taxa differentially distributed over time were further assessed by Dunn’s post-hoc test. The within-group homogeneity was tested by calculating multivariate dispersion indices with PRIMER.

## Acknowledgments

We thank Susanne Talay for her excellent introduction and support with all aspects of work in the BSL3 facilities at the Helmholtz Centre for Infection Research, the staff of the BSL3 animal facility for their help and support, Luiz Borges and Howard Junca for critical discussions and Ingo Schmitz for critically reading the manuscript. Petra Dersch and Sabrina Mühlen were supported by the German Centre for Infection Research (DZIF). We declare no conflicts of interest.

## Supplementary Information

### Supplementary Figures

**Fig. S1.**
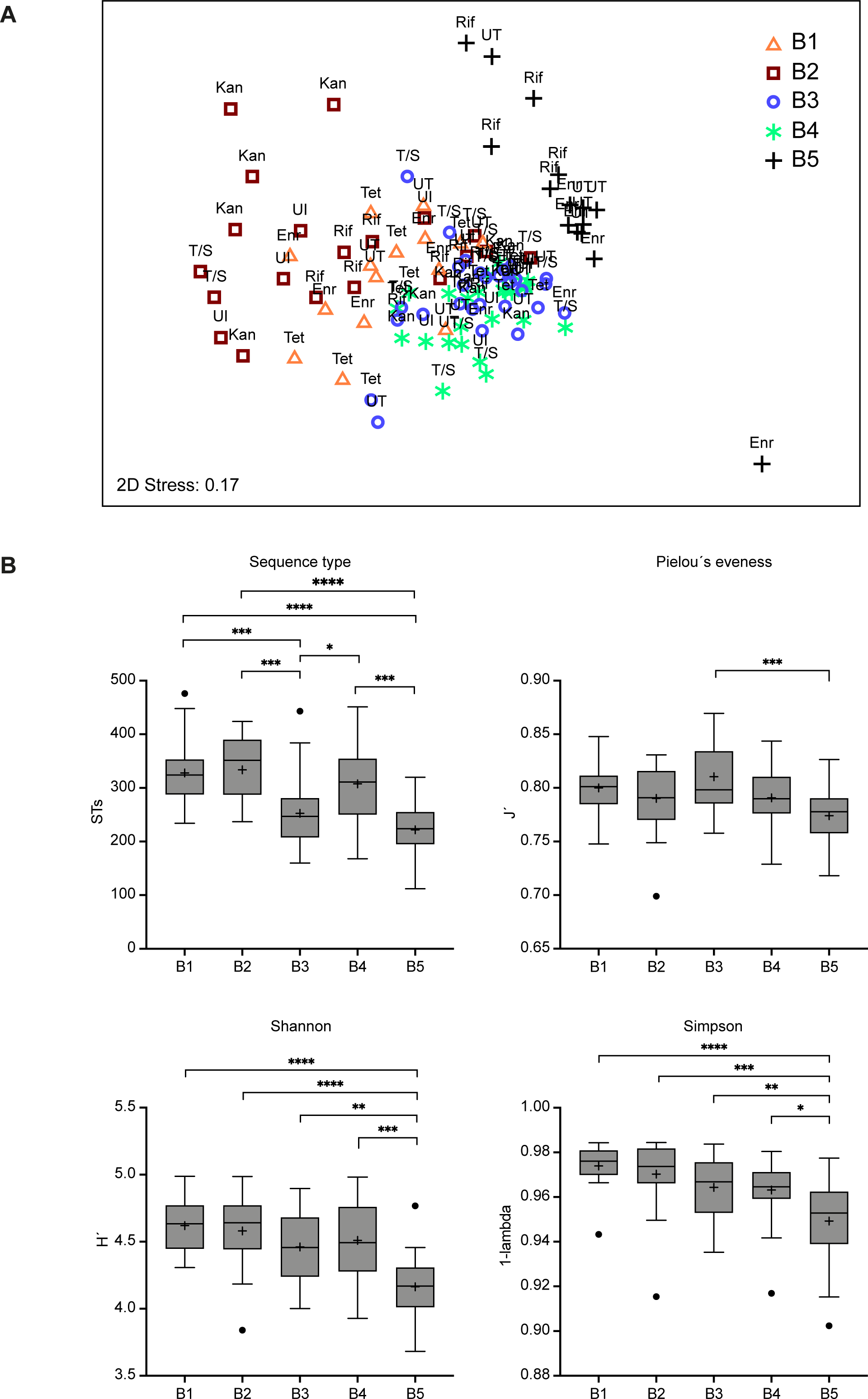
Global community structures of different mouse batches (experiments). **(A)** nMDS plot indicating the fecal global bacterial community structure of the separate mouse batches (indicated by different icons and colors) and their assignment to treatment groups (indicated above the icons; UI: uninfected, UT: untreated, Enf: Enrofloxacin, Kan: Kanamycin, Rif: Rifampicin, Tet: Tetracycline, T/S: Trimethoprim/Sulfamethoxazole). The global community structure is based on standardized sequence-type abundance data, and similarities were calculated using the Bray-Curtis similarity algorithm. **(B)** Diversity of the microbial communities in different mouse batches as indicated by total sequence-type number, Pielou’s evenness (J’) Shannon diversity (H’) and Simpsons diversity (1-λ), respectively, and was analyzed using sequence type relative abundance data as input. Data are based on an ordinary ANOVA analysis with Tukey’s test for multiple comparisons. Statistically significant differences are indicated as *p<0.05, **p<0.01, ***p<0.001 or ****p<0.0001. The mean is indicated by + and the median by a black line. The box represents the interquartile range. The whiskers extend to the upper adjacent value (largest value = 75th percentile + 1.5 x IQR) and the lower adjacent value (lowest value = 25th percentile - 1.5 x IQR) and the dots represent outliers.

**Fig. S2.**
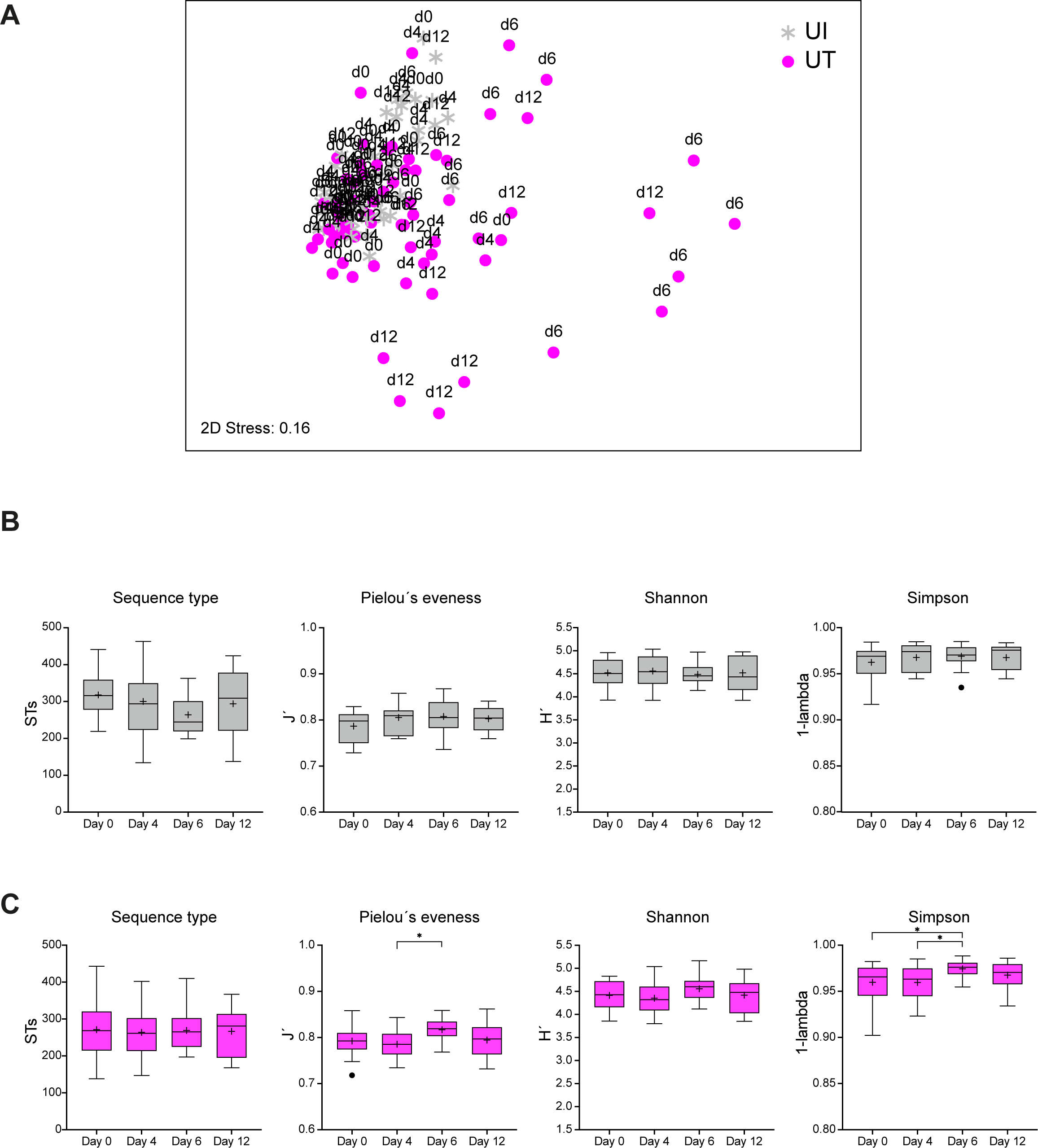
Global community structures of uninfected and infected untreated mice over time. (**A**) nMDS plot showing global bacterial community structures of UI (uninfected) and UT (untreated) mice over time. The time of sampling is indicated above the icons. The global community structure is based on standardized sequence type abundance data, and similarities were calculated using the Bray-Curtis similarity algorithm (**B** and **C**). Diversity of the microbial communities in UI (uninfected, **B**) and UT (untreated, **C**) mice as indicated by total sequence type number, Pielou’s evenness (J’), Shannon diversity (H’), and Simpsons diversity (1-λ), respectively, were analyzed using sequence type relative abundance data as input. Differences in diversity were tested by using a mixed effect model, and multiple comparisons were corrected using the Tukey test. Statistically significant differences are indicated as *p<0.05. The mean is indicated by + and the median by a black line. The box represents the interquartile range. The whiskers extend to the upper adjacent value (largest value = 75th percentile + 1.5 x IQR) and the lower adjacent value (lowest value = 25th percentile - 1.5 x IQR) and the dots represent outliers.

**Fig. S3.**
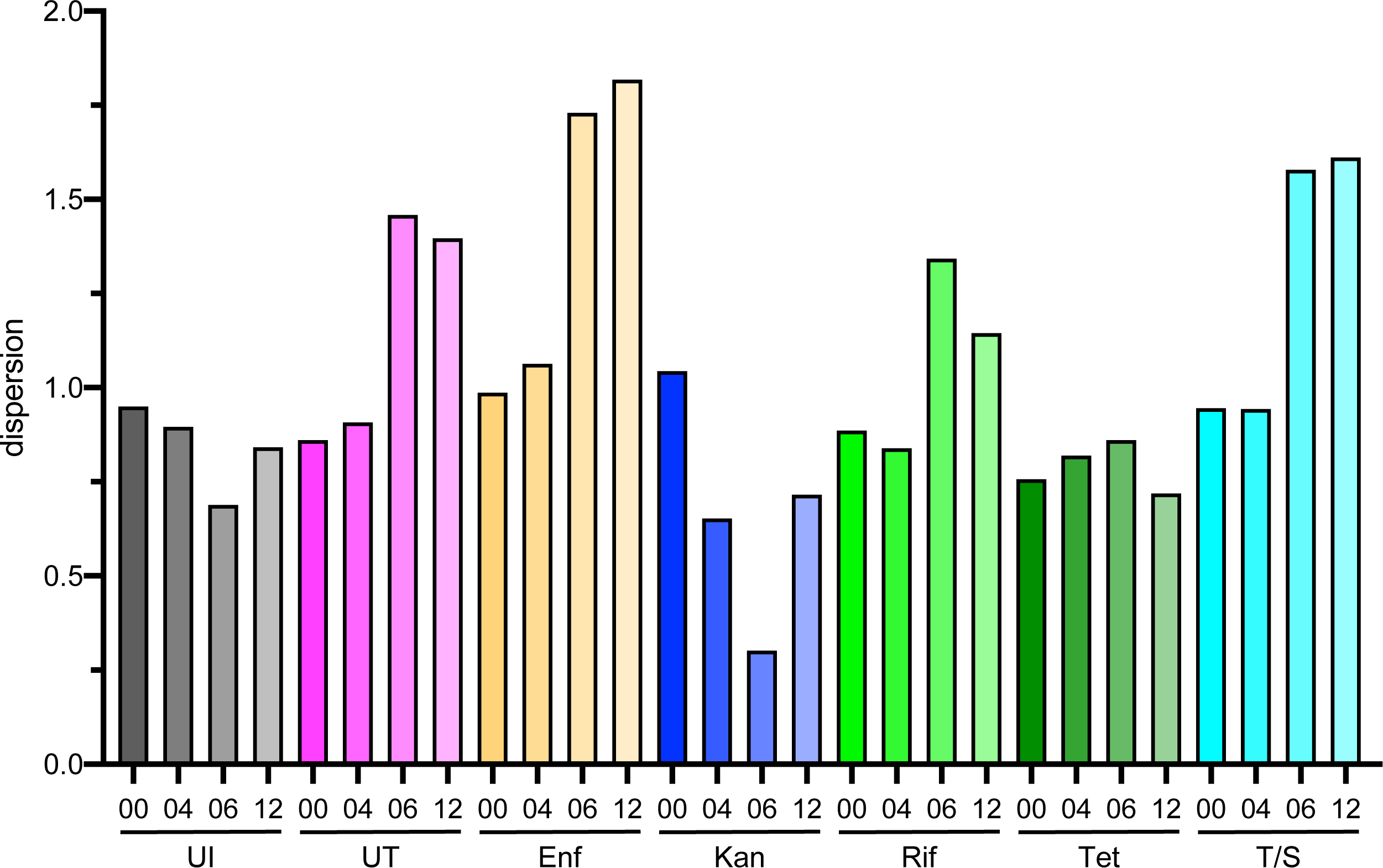
Heterogeneity in microbial communities depending on treatment and treatment times as indicated by multivariate dispersion indices. Indices indicating the within-group homogeneity were calculated by PRIMER and were followed for all treatment groups over time (UI, uninfected untreated; UT, infected untreated; Enf, Enrofloxacin; Kan, Kanamycin; Rif, Rifampicin; Tet, Tetracycline; T/S, Trimethoprim/Sulfamethoxazole).

**Fig. S4.**
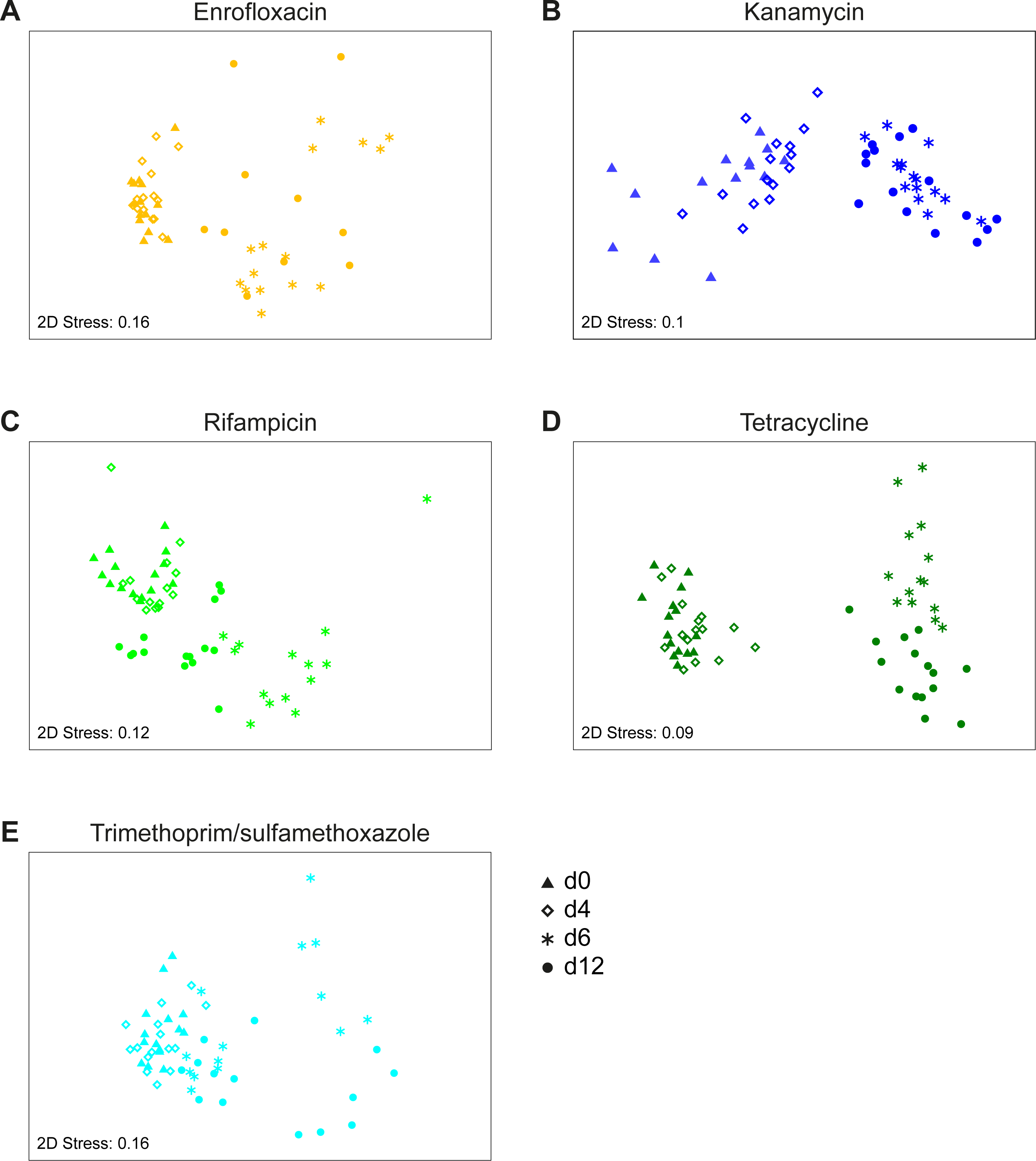
Relative abundance of *Bacteroides* species as influenced by infection and antibiotic treatment. Variations in the relative abundance of *Bacteroides*_11, *Bacteroides acidifaciens*, and *Bacteroides uniformis* were followed for all treatment groups over time (**A**, uninfected untreated; **B**, infected untreated; **C**, Enrofloxacin; **D**, Kanamycin; **E**, Rifampicin; **F**, Tetracycline; **G**, Trimethoprim/Sulfamethoxazole). Significant differences upon treatment time were calculated using the Kruskal-Wallis test with Benjamini-Hochberg correction and are indicated as **p<0.01, ***p<0.001 and ****p<0.0001. The individual relative abundances are given, and the mean (dotted lines) ± SEM is indicated.

**Fig. S5.**
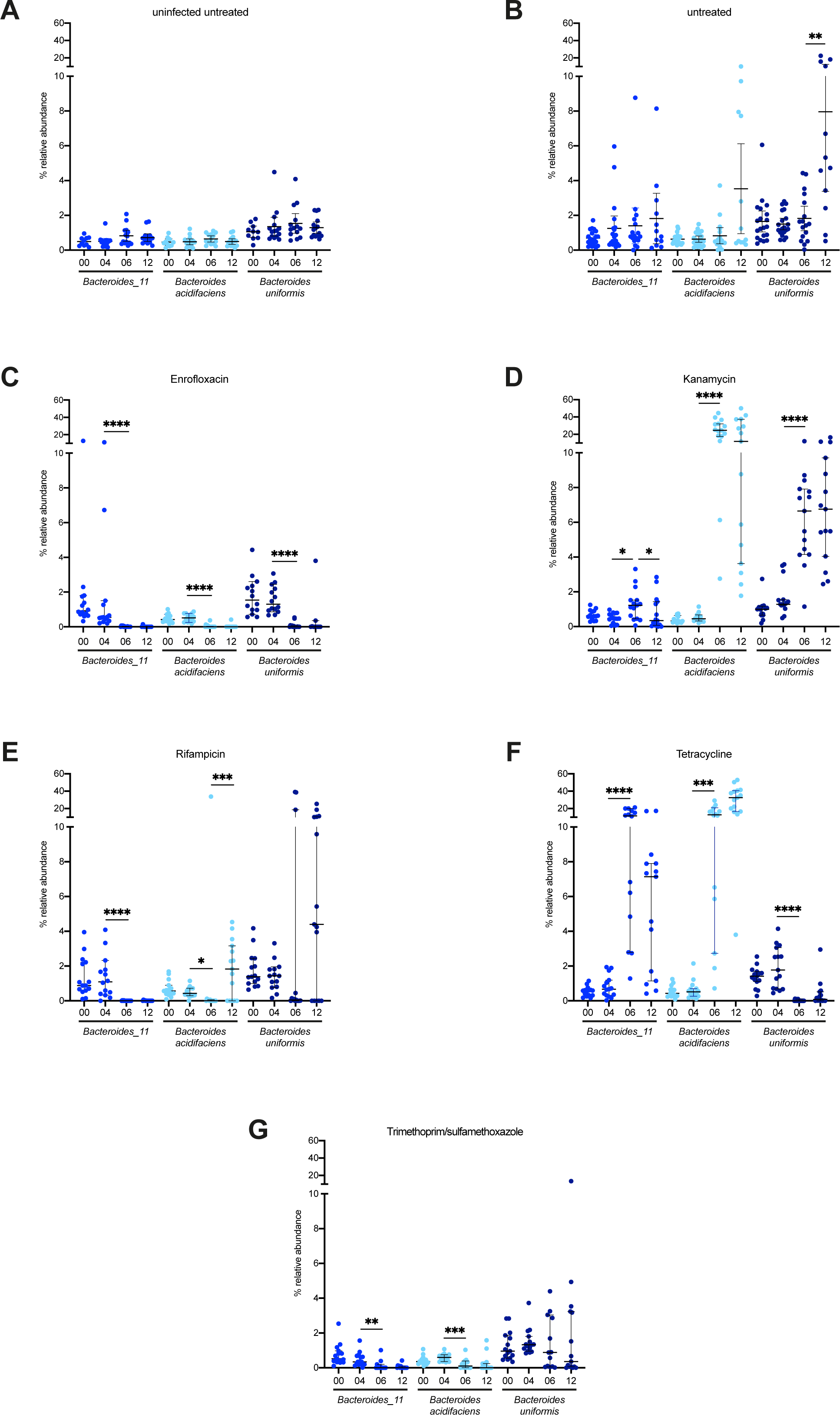
Differences in global bacterial community structure in mice feces upon infection and subsequent antibiotic treatment. The global bacterial community structure was assessed by non-metric multidimensional scaling (nMDS) and is based on standardized sequence type abundance data. Similarities were calculated using the Bray-Curtis similarity algorithm. All treatment groups were infected with *C. rodentium* ϕ*stx2_dact_* on day 0. Treatment groups received antibiotics (**A**) enrofloxacin, (**B**) kanamycin, (**C**) rifampicin, (**D**) tetracycline, or (**E**) trimethoprim/sulfamethoxazole from day 4 post-infection.

**Fig. S6.**
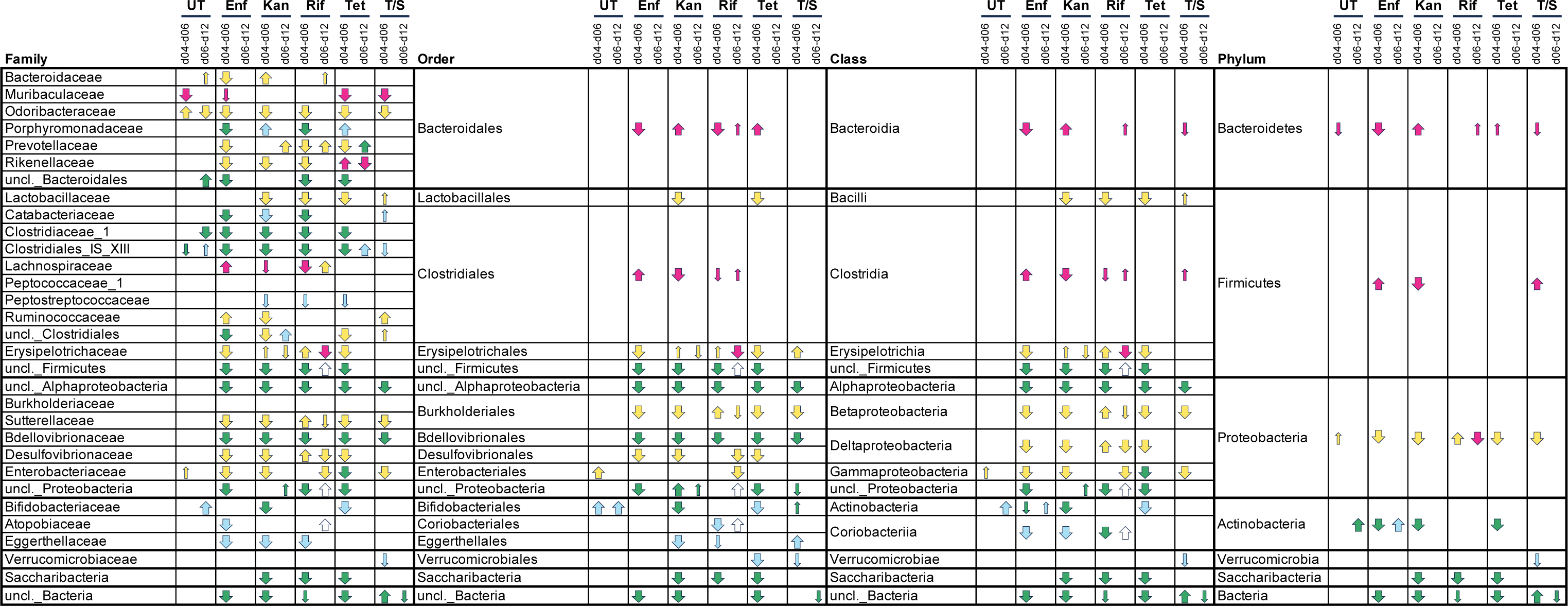
Phylogenetic taxa (families, order, classes, and phyla) influenced by infection and antibiotic treatment. The abundances of taxa were compared using the Kruskal-Wallis test with Benjamini-Hochberg corrections for multiple comparisons. Groups of samples were considered significantly different in abundance if the adjusted p-value was <0.05. Taxa differentially distributed over time were further assessed by Dunn’s post-hoc test. Changes in relative abundance between day 4 (d04) and day 6 (d06) and between day 6 (d06) and day 12 (d12) during infection (UT) and antibiotic treatment (Enf, enrofloxacin; Kan, kanamycin; Rif, rifampicin; Tet, tetracycline; or T/S, trimethoprim/ sulfamethoxazole) are indicated by arrows with the arrow direction indicating increase or decrease in abundance. A bold arrow indicates a significant change in relative abundance with p<0.01 and a small arrow with 0.01<p<0.05. The color of the arrow shows the relative abundance on d4 or d6 (red: >10%, yellow: 1-10%; green 0.1-1%, blue <0.1%).

**Fig. S7.**
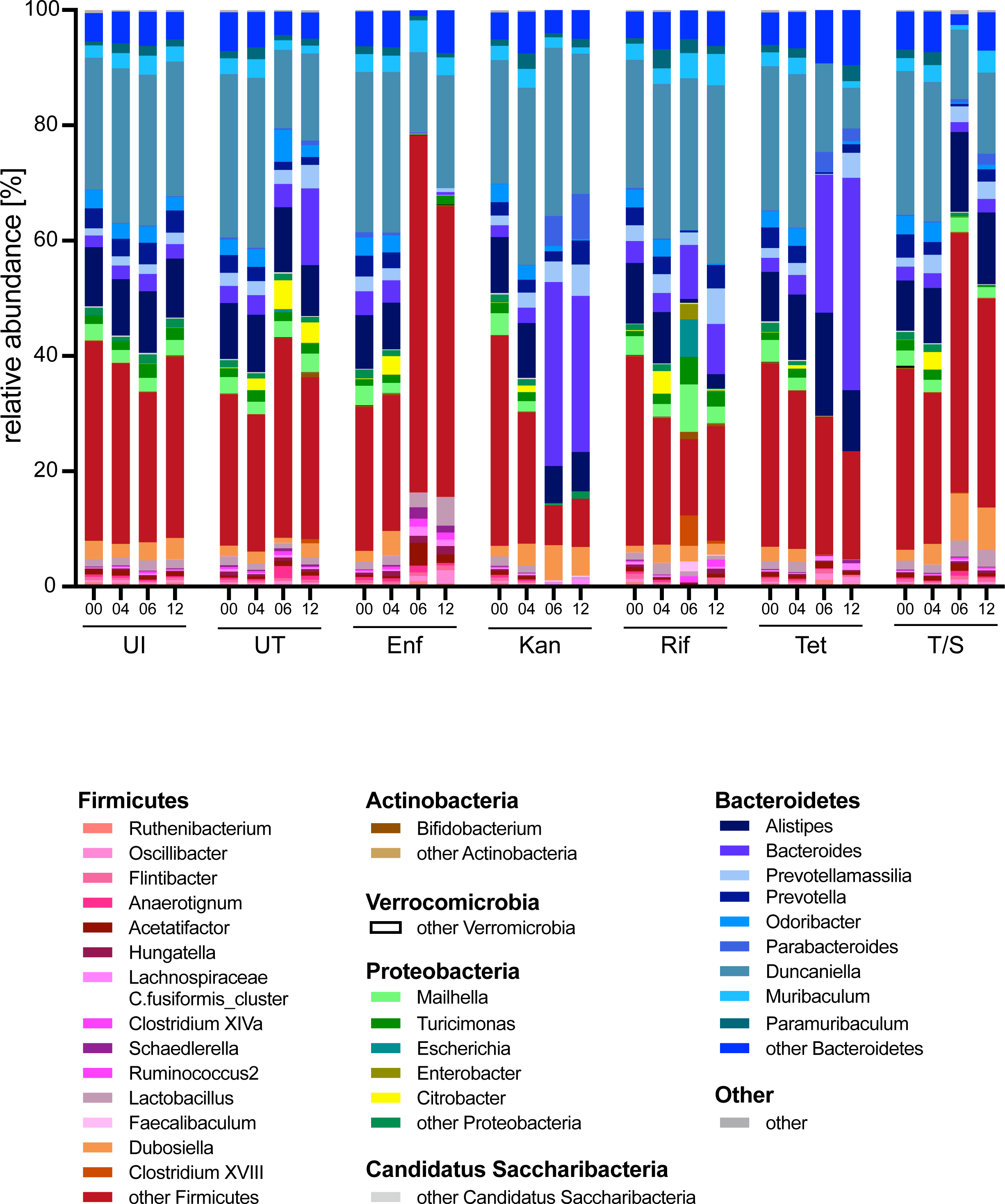
Stacked bar-plot representation of microbiota compositions at the genus level. The relative abundance of the different genera in the fecal microbiota of uninfected mice, infected mice, and mice additionally treated with antibiotics is given for all time points assessed. Genera with an abundance below 1% as well as taxa that could not be classified down to the genus level but at least to the phyla given are summarized to the respective phylum. Sequences that could not be classified to the phylum level are grouped as other. The different treatment groups (uninfected untreated (UI); infected untreated (UT); Enf, enrofloxacin; Kan, kanamycin; Rif, rifampicin; Tet, tetracycline; or T/S, trimethoprim/ sulfamethoxazole) are given below the X-axis label indicating the sampling day (00: D0; 04: D4; 06: D6; and 12: D12).

### Supplementary Tables

**Table S1: Nucleotide sequences of all sequence variants determined using Illumina-based amplicon deep-sequencing, their phylogenetic annotation and sequence count as well as relative abundance data after rarefying across all 410 fecal samples.**

**Table S2:** Factors influencing global community structures prior to infection as indicated by PERMANOVA. The significance of differences in community structure between different mouse batches (experiments) and treatment groups was calculated by PERMANOVA (main test). The Pseudo-F and the Monte Carlo p-values are given for each factor performed at different taxonomic levels (from sequence type to phylum). The t statistics and the Monte Carlo p-values are also given for paired tests among different mouse batches. Analysis was performed at different taxonomic levels (from sequence type to phylum). Bold p <0.05.

| Factor (main test) | Sequence type |  | Genus |  | Family |  | Order |  | Class |  | Phylum |  |
| --- | --- | --- | --- | --- | --- | --- | --- | --- | --- | --- | --- | --- |
|  | Pseudo-F | p(MC) | Pseudo-F | p(MC) | Pseudo-F | p(MC) | Pseudo-F | p(MC) | Pseudo-F | p(MC) | Pseudo-F | p(MC) |
| Mouse batch | 7.65 | **0.001** | 9.892 | **0.001** | 9.1471 | **0.001** | 7.0709 | **0.001** | 7.1082 | **0.001** | 5.5688 | **0.001** |
| Treatment | 1.265<br>4 | 0.05 | 0.9964 | 0.444 | 0.9504 | 0.486 | 0.8906 | 0.500 | 0.8832 | 0.528 | 0.8772 | 0.522 |

| Mouse batch | t | p(MC) | t | p(MC) | t | p(MC) | t | p(MC) | t | p(MC) | t | p(MC) |
| --- | --- | --- | --- | --- | --- | --- | --- | --- | --- | --- | --- | --- |
| Exp1, Exp2 | 1.4338 | **0.03** | 1.5872 | 0.069 | 1.6066 | 0.086 | 1.4597 | 0.152 | 1.4548 | 0.145 | 1.4577 | 0.154 |
| Exp1, Exp3 | 2.5183 | **0.001** | 3.1116 | **0.001** | 3.2787 | **0.001** | 2.6604 | **0.005** | 2.6904 | **0.008** | 2.414 | **0.019** |
| Exp1, Exp4 | 2.5919 | **0.001** | 2.9979 | **0.001** | 3.102 | **0.001** | 2.2253 | **0.021** | 2.2552 | **0.03** | 0.9391 | 0.361 |
| Exp1, Exp5 | 3.0989 | **0.001** | 3.2492 | **0.001** | 2.6754 | **0.002** | 2.9616 | **0.007** | 2.986 | **0.003** | 2.9594 | **0.003** |
| Exp2, Exp3 | 3.0072 | **0.001** | 4.531 | **0.001** | 4.6244 | **0.001** | 4.1387 | **0.001** | 4.146 | **0.001** | 3.8986 | **0.001** |
| Exp2, Exp4 | 3.1034 | **0.001** | 4.3015 | **0.001** | 4.3475 | **0.001** | 3.5471 | **0.001** | 3.5569 | **0.002** | 2.6147 | **0.016** |
| Exp2, Exp5 | 3.3833 | **0.001** | 4.1237 | **0.001** | 3.6481 | **0.001** | 3.8001 | **0.001** | 3.7982 | **0.002** | 3.8884 | **0.002** |
| Exp3, Exp4 | 1.9542 | **0.001** | 1.5691 | 0.077 | 1.5049 | 0.089 | 1.5351 | 0.123 | 1.5206 | 0.11 | 1.8825 | 0.056 |
| Exp3, Exp5 | 3.2861 | **0.001** | 2.5251 | **0.001** | 1.9199 | 0.021 | 1.4215 | 0.156 | 1.3729 | 0.152 | 1.0841 | 0.296 |
| Exp4, Exp5 | 3.477 | **0.001** | 3.1231 | **0.001** | 2.4874 | 0.006 | 2.4307 | **0.007** | 2.425 | **0.012** | 2.6552 | **0.012** |

**Table S3:**
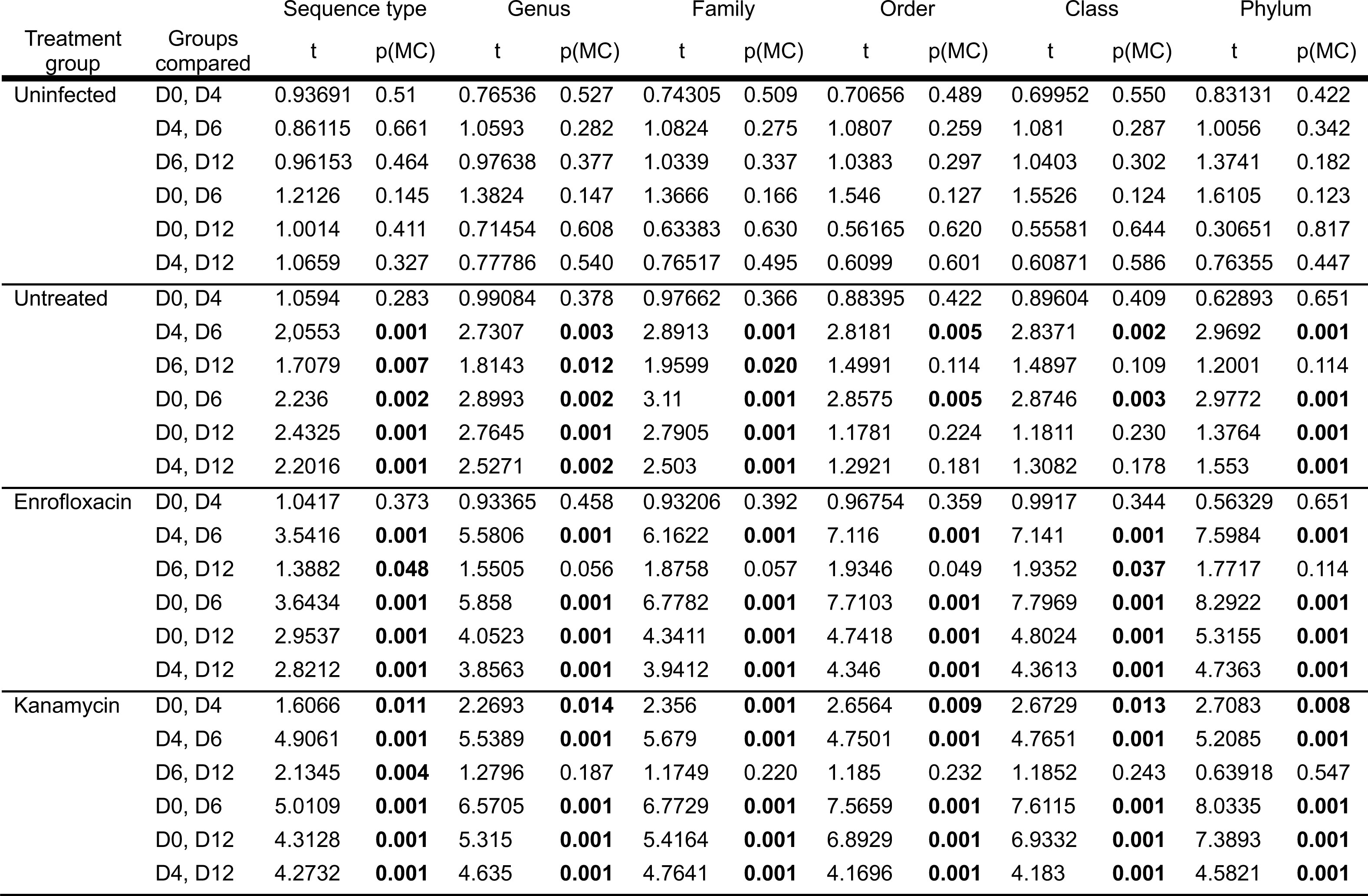
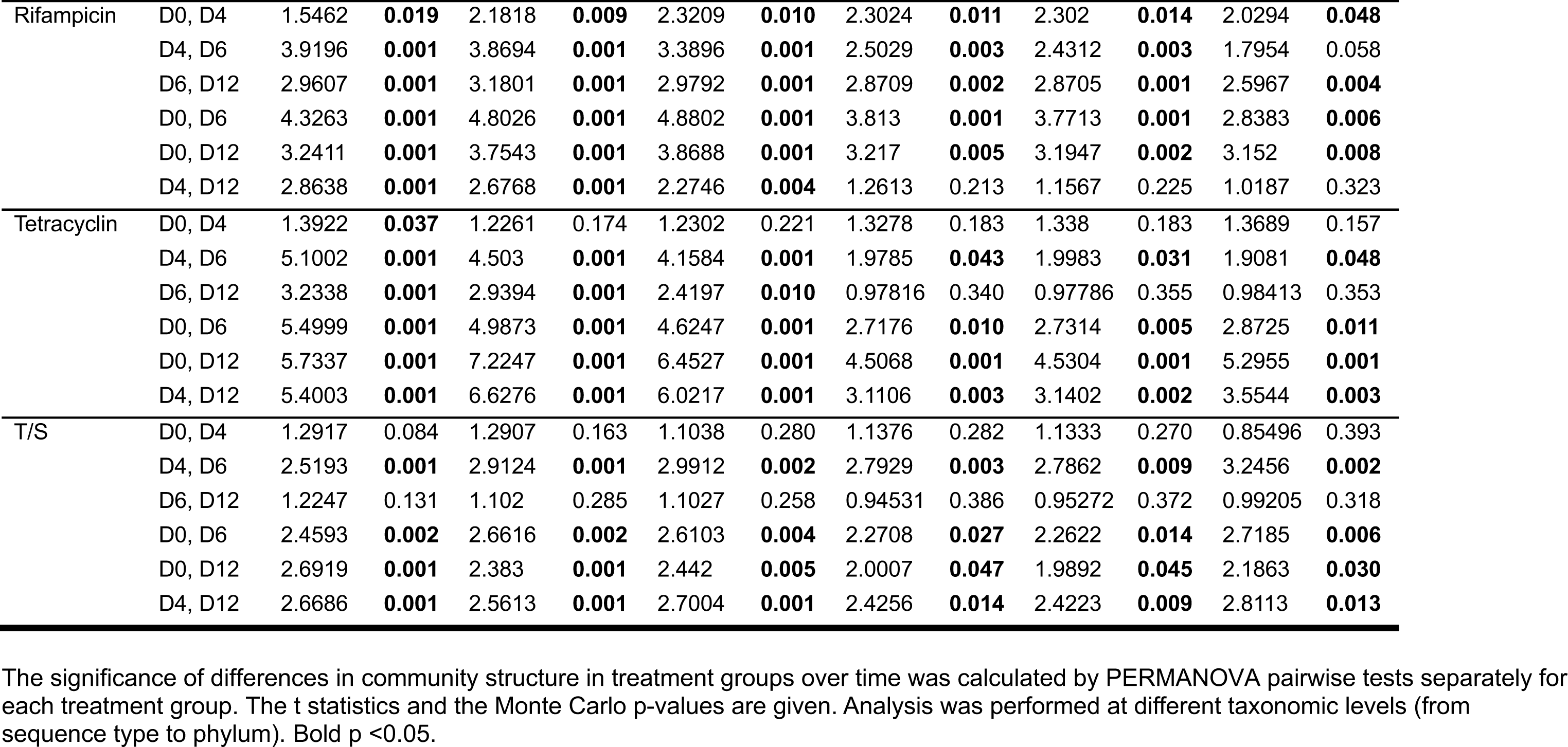
Influence of infection and antibiotic treatment on the community structure as indicated by PERMANOVA. The significance of differences in community structure in treatment groups over time was calculated by PERMANOVA pairwise tests separately for each treatment group. The t statistics and the Monte Carlo p-values are given. Analysis was performed at different taxonomic levels (from sequence type to phylum). Bold p <0.05.

| Treatment group | Groups compared | Sequence type |  | Genus |  | Family |  | Order |  | Class |  | Phylum |  |
| --- | --- | --- | --- | --- | --- | --- | --- | --- | --- | --- | --- | --- | --- |
|  |  | t | p(MC) | t | p(MC) | t | p(MC) | t | p(MC) | t | p(MC) | t | p(MC) |
| Uninfected | D0, D4 | 0.93691 | 0.51 | 0.76536 | 0.527 | 0.74305 | 0.509 | 0.70656 | 0.489 | 0.69952 | 0.550 | 0.83131 | 0.422 |
|  | D4, D6 | 0.86115 | 0.661 | 1.0593 | 0.282 | 1.0824 | 0.275 | 1.0807 | 0.259 | 1.081 | 0.287 | 1.0056 | 0.342 |
|  | D6, D12 | 0.96153 | 0.464 | 0.97638 | 0.377 | 1.0339 | 0.337 | 1.0383 | 0.297 | 1.0403 | 0.302 | 1.3741 | 0.182 |
|  | D0, D6 | 1.2126 | 0.145 | 1.3824 | 0.147 | 1.3666 | 0.166 | 1.546 | 0.127 | 1.5526 | 0.124 | 1.6105 | 0.123 |
|  | D0, D12 | 1.0014 | 0.411 | 0.71454 | 0.608 | 0.63383 | 0.630 | 0.56165 | 0.620 | 0.55581 | 0.644 | 0.30651 | 0.817 |
|  | D4, D12 | 1.0659 | 0.327 | 0.77786 | 0.540 | 0.76517 | 0.495 | 0.6099 | 0.601 | 0.60871 | 0.586 | 0.76355 | 0.447 |
| Untreated | D0, D4 | 1.0594 | 0.283 | 0.99084 | 0.378 | 0.97662 | 0.366 | 0.88395 | 0.422 | 0.89604 | 0.409 | 0.62893 | 0.651 |
|  | D4, D6 | 2.0553 | **0.001** | 2.7307 | **0.003** | 2.8913 | **0.001** | 2.8181 | **0.005** | 2.8371 | **0.002** | 2.9692 | **0.001** |
|  | D6, D12 | 1.7079 | **0.007** | 1.8143 | **0.012** | 1.9599 | **0.020** | 1.4991 | 0.114 | 1.4897 | 0.109 | 1.2001 | 0.114 |
|  | D0, D6 | 2.236 | **0.002** | 2.8993 | **0.002** | 3.11 | **0.001** | 2.8575 | **0.005** | 2.8746 | **0.003** | 2.9772 | **0.001** |
|  | D0, D12 | 2.4325 | **0.001** | 2.7645 | **0.001** | 2.7905 | **0.001** | 1.1781 | 0.224 | 1.1811 | 0.230 | 1.3764 | **0.001** |
|  | D4, D12 | 2.2016 | **0.001** | 2.5271 | **0.002** | 2.503 | **0.001** | 1.2921 | 0.181 | 1.3082 | 0.178 | 1.553 | **0.001** |
| Enrofloxacin | D0, D4 | 1.0417 | 0.373 | 0.93365 | 0.458 | 0.93206 | 0.392 | 0.96754 | 0.359 | 0.9917 | 0.344 | 0.56329 | 0.651 |
|  | D4, D6 | 3.5416 | **0.001** | 5.5806 | **0.001** | 6.1622 | **0.001** | 7.116 | **0.001** | 7.141 | **0.001** | 7.5984 | **0.001** |
|  | D6, D12 | 1.3882 | **0.048** | 1.5505 | 0.056 | 1.8758 | 0.057 | 1.9346 | 0.049 | 1.9352 | **0.037** | 1.7717 | 0.114 |
|  | D0, D6 | 3.6434 | **0.001** | 5.858 | **0.001** | 6.7782 | **0.001** | 7.7103 | **0.001** | 7.7969 | **0.001** | 8.2922 | **0.001** |
|  | D0, D12 | 2.9537 | **0.001** | 4.0523 | **0.001** | 4.3411 | **0.001** | 4.7418 | **0.001** | 4.8024 | **0.001** | 5.3155 | **0.001** |
|  | D4, D12 | 2.8212 | **0.001** | 3.8563 | **0.001** | 3.9412 | **0.001** | 4.346 | **0.001** | 4.3613 | **0.001** | 4.7363 | **0.001** |
| Kanamycin | D0, D4 | 1.6066 | **0.011** | 2.2693 | **0.014** | 2.356 | **0.001** | 2.6564 | **0.009** | 2.6729 | **0.013** | 2.7083 | **0.008** |
|  | D4, D6 | 4.9061 | **0.001** | 5.5389 | **0.001** | 5.679 | **0.001** | 4.7501 | **0.001** | 4.7651 | **0.001** | 5.2085 | **0.001** |
|  | D6, D12 | 2.1345 | **0.004** | 1.2796 | 0.187 | 1.1749 | 0.220 | 1.185 | 0.232 | 1.1852 | 0.243 | 0.63918 | 0.547 |
|  | D0, D6 | 5.0109 | **0.001** | 6.5705 | **0.001** | 6.7729 | **0.001** | 7.5659 | **0.001** | 7.6115 | **0.001** | 8.0335 | **0.001** |
|  | D0, D12 | 4.3128 | **0.001** | 5.315 | **0.001** | 5.4164 | **0.001** | 6.8929 | **0.001** | 6.9332 | **0.001** | 7.3893 | **0.001** |
|  | D4, D12 | 4.2732 | **0.001** | 4.635 | **0.001** | 4.7641 | **0.001** | 4.1696 | **0.001** | 4.183 | **0.001** | 4.5821 | **0.001** |
| Rifampicin | D0, D4 | 1.5462 | **0.019** | 2.1818 | **0.009** | 2.3209 | **0.010** | 2.3024 | **0.011** | 2.302 | **0.014** | 2.0294 | **0.048** |
|  | D4, D6 | 3.9196 | **0.001** | 3.8694 | **0.001** | 3.3896 | **0.001** | 2.5029 | **0.003** | 2.4312 | **0.003** | 1.7954 | 0.058 |
|  | D6, D12 | 2.9607 | **0.001** | 3.1801 | **0.001** | 2.9792 | **0.001** | 2.8709 | **0.002** | 2.8705 | **0.001** | 2.5967 | **0.004** |
|  | D0, D6 | 4.3263 | **0.001** | 4.8026 | **0.001** | 4.8802 | **0.001** | 3.813 | **0.001** | 3.7713 | **0.001** | 2.8383 | **0.006** |
|  | D0, D12 | 3.2411 | **0.001** | 3.7543 | **0.001** | 3.8688 | **0.001** | 3.217 | **0.005** | 3.1947 | **0.002** | 3.152 | **0.008** |
|  | D4, D12 | 2.8638 | **0.001** | 2.6768 | **0.001** | 2.2746 | **0.004** | 1.2613 | 0.213 | 1.1567 | 0.225 | 1.0187 | 0.323 |
| Tetracyclin | D0, D4 | 1.3922 | **0.037** | 1.2261 | 0.174 | 1.2302 | 0.221 | 1.3278 | 0.183 | 1.338 | 0.183 | 1.3689 | 0.157 |
|  | D4, D6 | 5.1002 | **0.001** | 4.503 | **0.001** | 4.1584 | **0.001** | 1.9785 | **0.043** | 1.9983 | **0.031** | 1.9081 | **0.048** |
|  | D6, D12 | 3.2338 | **0.001** | 2.9394 | **0.001** | 2.4197 | **0.010** | 0.97816 | 0.340 | 0.97786 | 0.355 | 0.98413 | 0.353 |
|  | D0, D6 | 5.4999 | **0.001** | 4.9873 | **0.001** | 4.6247 | **0.001** | 2.7176 | **0.010** | 2.7314 | **0.005** | 2.8725 | **0.011** |
|  | D0, D12 | 5.7337 | **0.001** | 7.2247 | **0.001** | 6.4527 | **0.001** | 4.5068 | **0.001** | 4.5304 | **0.001** | 5.2955 | **0.001** |
|  | D4, D12 | 5.4003 | **0.001** | 6.6276 | **0.001** | 6.0217 | **0.001** | 3.1106 | **0.003** | 3.1402 | **0.002** | 3.5544 | **0.003** |
| T/S | D0, D4 | 1.2917 | 0.084 | 1.2907 | 0.163 | 1.1038 | 0.280 | 1.1376 | 0.282 | 1.1333 | 0.270 | 0.85496 | 0.393 |
|  | D4, D6 | 2.5193 | **0.001** | 2.9124 | **0.001** | 2.9912 | **0.002** | 2.7929 | **0.003** | 2.7862 | **0.009** | 3.2456 | **0.002** |
|  | D6, D12 | 1.2247 | 0.131 | 1.102 | 0.285 | 1.1027 | 0.258 | 0.94531 | 0.386 | 0.95272 | 0.372 | 0.99205 | 0.318 |
|  | D0, D6 | 2.4593 | **0.002** | 2.6616 | **0.002** | 2.6103 | **0.004** | 2.2708 | **0.027** | 2.2622 | **0.014** | 2.7185 | **0.006** |
|  | D0, D12 | 2.6919 | **0.001** | 2.383 | **0.001** | 2.442 | **0.005** | 2.0007 | **0.047** | 1.9892 | **0.045** | 2.1863 | **0.030** |
|  | D4, D12 | 2.6686 | **0.001** | 2.5613 | **0.001** | 2.7004 | **0.001** | 2.4256 | **0.014** | 2.4223 | **0.009** | 2.8113 | **0.013** |

**Table S4: Phylogenetic taxa (genera, families, order, classes, and phyla) influenced by infection and antibiotic treatment.** Changes over time were assessed by the Kruskal-Wallis test with Benjamini-Hochberg corrections for multiple comparisons separately in uninfected mice (UI), infected but untreated mice (UT) as well as in Enf (enrofloxacin), Kan (kanamycin), Rif (rifampicin), Tet (tetracycline), or T/S (trimethoprim/sulfamethoxazole) treated mice. Groups of samples were considered significantly different if the adjusted p-value was <0.05. Both the original (KW_p) as well as the adjusted p-value (KW_padj_BH) are given. Taxa differentially distributed over time were further assessed by Dunn’s post-hoc test. The significance of the taxon difference between the different time points is given. p-values >0.01 are indicated in yellow and p-values between 0.05 and 0.01 in orange.

**Table S5:**
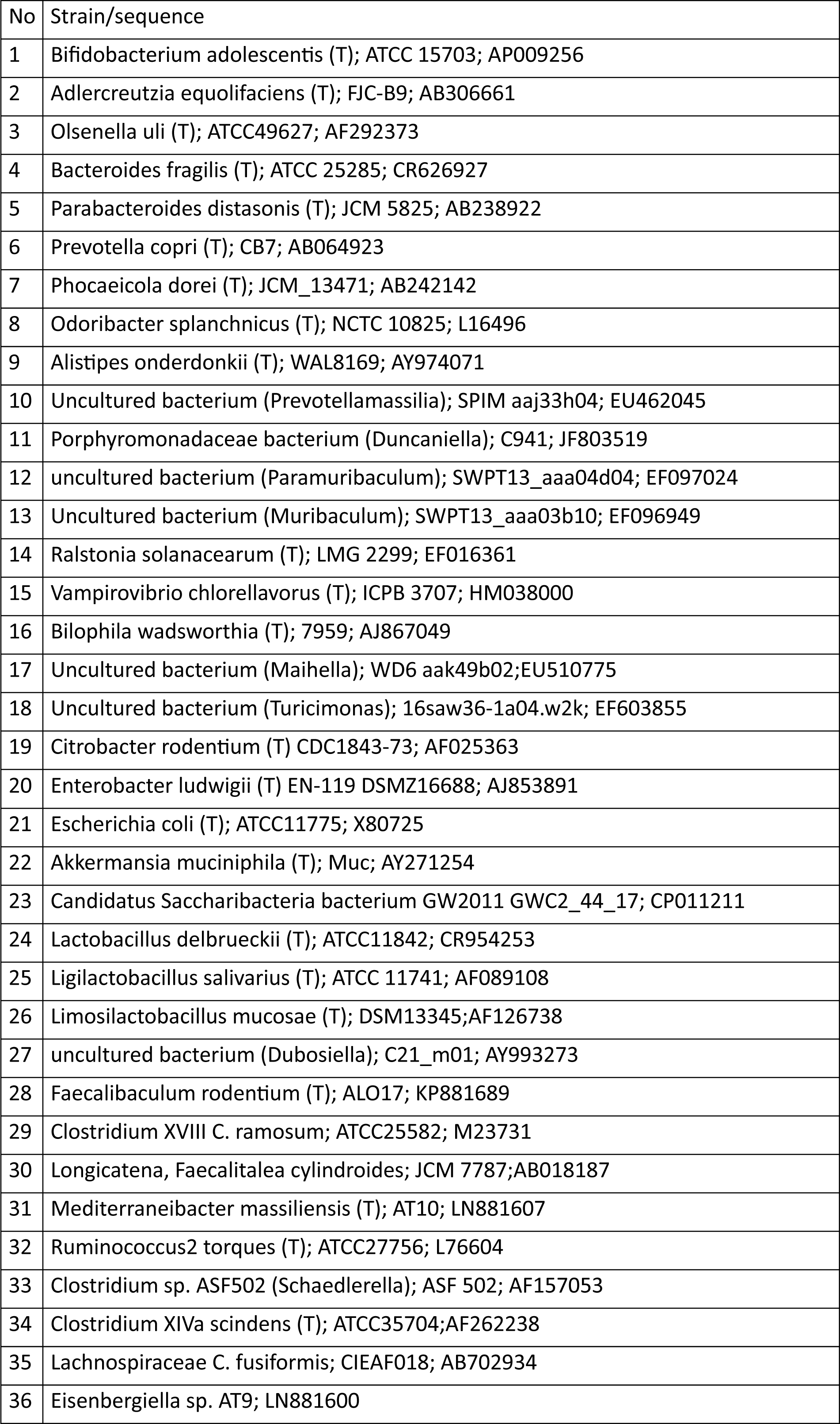
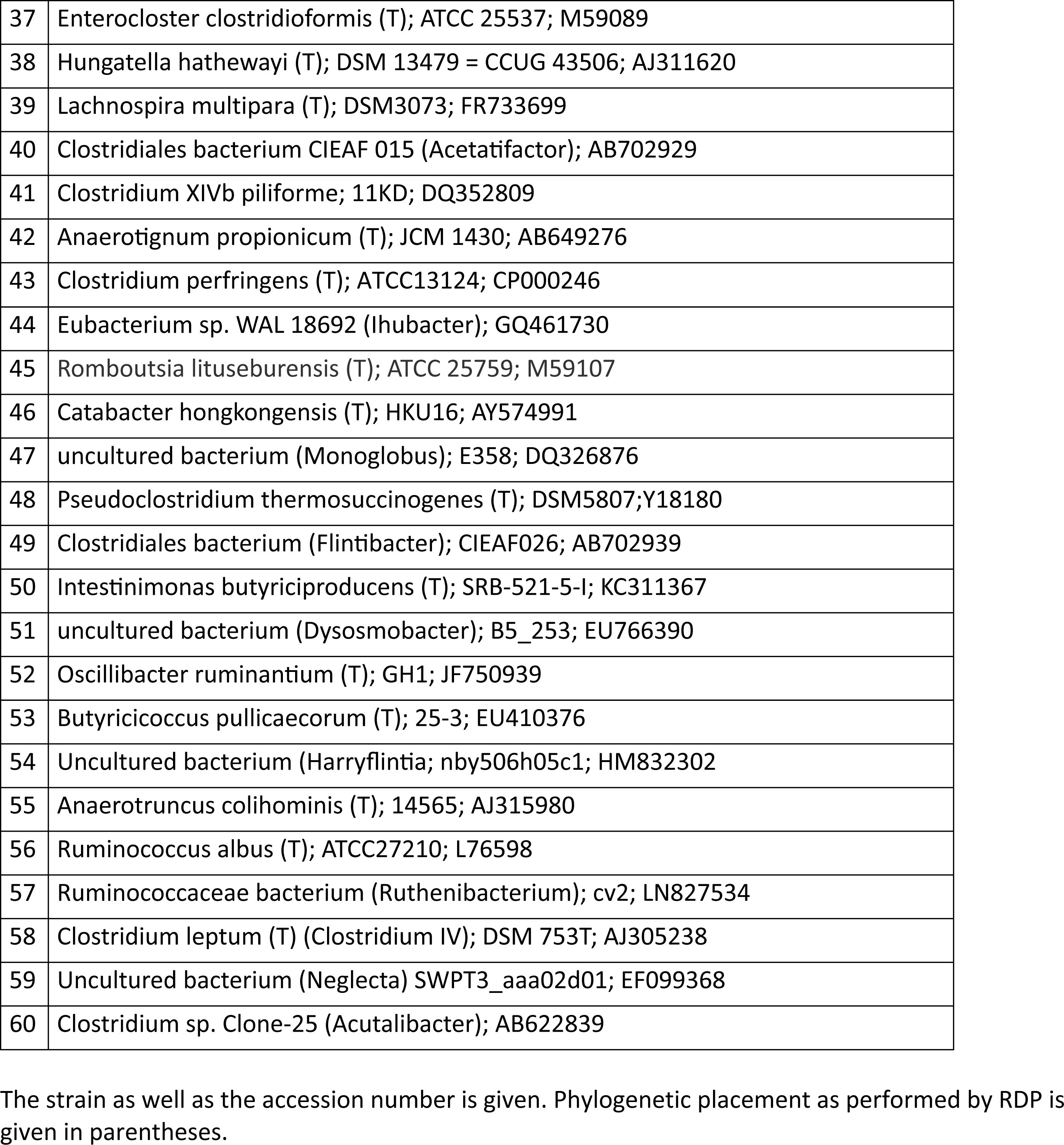
Sequences used as representative for genera identified. The strain as well as the accession number is given. Phylogenetic placement as performed by RDP is given in parentheses.

